# Air pollution induces *Staphylococcus aureus* USA300 respiratory tract colonisation mediated by specific bacterial genetic responses dependent on the global virulence gene regulators Agr and Sae

**DOI:** 10.1101/2022.02.04.479102

**Authors:** Jo Purves, Shane. J. K. Hussey, Louise Corscadden, Lillie Purser, Andie Hall, Raju Misra, Paul S. Monks, Julian M. Ketley, Peter W. Andrew, Julie A. Morrissey

**Author notes:** Corresponding Author: Prof. Julie. A. Morrissey. Postal Address: Prof. Julie Morrissey, Department of Genetics and Genome Biology, University of Leicester, Leicester, LE1 7RH.

## Abstract

Exposure to particulate matter (PM), a major component of air pollution, is associated with exacerbation of chronic respiratory disease, and infectious diseases such as community acquired pneumonia. Although PM can cause adverse health effects through direct damage to host cells, our previous study showed that PM can also impact bacterial behaviour by promoting *in vivo* colonisation. In this study we describe the genetic mechanisms involved in the bacterial response to exposure to black carbon (BC), a constituent of PM found in most sources of air pollution. We show that *Staphylococcus aureus* strain USA300 LAC grown in BC prior to inoculation showed increased murine respiratory tract colonisation and pulmonary invasion *in vivo*, as well as adhesion and invasion of human epithelial cells *in vitro*. Global transcriptional analysis showed that BC has a widespread effect on *S. aureus* transcriptional responses, altering the regulation of the major virulence gene regulators Sae and Agr and causing increased expression of genes encoding toxins, proteases, and immune evasion factors. Together these data describe a previously unrecognised causative mechanism of air pollution-associated infection, in that exposure to BC can increase bacterial colonisation and virulence factor expression by acting directly on the bacterium rather than via the host.

**Originality-Significance Statement:** This study shows that exposure to air pollution results in a global change in gene expression in bacteria. Specifically, our data show that in the important human pathogen *Staphylococcus aureus*, exposure to a major constituent of air pollution, black carbon (BC) results in widespread changes in global gene expression, altering the expression of key virulence determinants. Furthermore, *S. aureus* that are exposed to BC prior to inoculation show increased colonisation of the murine nasopharynx and lungs *in vivo*, and increased adhesion and invasion in lung epithelial cells *in vitro*. These findings indicate that air pollution has a significant and direct impact on bacteria, altering their behaviour and their potential to colonise and invade during infection. While many studies have taken a host-focussed approach to studying the impact of air pollution on human health, this study takes a pathogen-focussed approach to further the understanding of these fundamental interactions to identify new causative mechanisms of the detrimental effects of air pollution. This is critical for understanding the adverse health effects caused by exposure to air pollution, the single largest environmental risk to human health in the world.

## Introduction

Air pollution is the world’s largest single global environmental health risk with an estimated 90% of people worldwide breathing polluted air, this pollution is responsible for over 7 million deaths per year (World Health Organization, - News Release, 2018). It is the result of natural and anthropogenic activity, with increased urbanisation resulting in significant increases in types and concentrations of pollutants (Manisalidis et al., 2020). Particulate matter (PM) is a major component of air pollution, with particles of <2.5 μm causing the most serious adverse health effects due to deposition in the upper respiratory tract and the ability to enter the lower respiratory tract and bloodstream (Cohen et al., 2017; McNeil, 2019).

PM exposure is strongly associated with cancer and cardiovascular diseases, and exacerbation of chronic respiratory disease, such as COPD and asthma (Cohen et al., 2017). There is also an association with infectious disease, with community acquired pneumonia rates most affected (Neupane et al., 2010; Qiu et al., 2014), but less well known is the impact on infective endocarditis (Hsieh et al., 2019), infection of cystic fibrosis patients (Psoter et al., 2015; Psoter et al., 2017), otitis media (Park et al., 2018), chronic rhinosinusitis (Schwarzbach et al., 2020) and adverse effects on chronic skin diseases (Dijkhoff et al., 2020). High levels of PM exposure also alter respiratory microbiome diversity (Li, R. et al., 2017; Li, X. et al., 2019; Mariani et al., 2018; Mariani et al., 2020; Rylance et al., 2016; Wang et al., 2019).

In seeking to explain how PM adversely affects chronic and infectious diseases, research has focused on direct damage to host tissue caused by PM exposure, including increased inflammation, and oxidative stress (Lee et al., 2021). It is also known that in infection, PM can potentiate disease by repressing the immune system (Castranova et al., 2001; Liu, J. et al., 2019; Migliaccio et al., 2013; Shears et al., 2020; Yang et al., 2001) and by disruption of epithelial function (Liu et al., 2019; Misiukiewicz-Stepien & Paplinska-Goryca, 2021). The possibility that PM may directly affect bacteria had not received attention until recently. In Hussey *et al*. (2017) we showed that air pollution does have a direct impact on bacterial behaviour.

Direct exposure of *Staphylococcus aureus* and *Streptococcus pneumoniae* to black carbon (BC), a by-product of biomass burning and a major constituent of PM (Bell et al., 2007), results in major changes in bacterial biofilm formation and antibiotic susceptibility (Hussey et al., 2017). Additionally, we found that in mice simultaneously exposed to *S. pneumoniae* and BC there was increased bacterial dissemination to the lungs (Hussey et al., 2017). The instillation of BC into the mice did not cause detectable tissue damage, indicating that BC acts as a signal that alters bacterial behaviour (Hussey et al., 2017). Subsequent studies also showed that direct bacterial exposure to a variety of PM sources also increased biofilm formation (Woo et al., 2018; Yadav et al., 2020) and increased *S. pneumoniae* nasopharyngeal colonisation and dissemination to the lungs and middle ear of mice (Yadav et al., 2020). None of these studies sought to determine the biological mechanisms involved in the bacterial response to PM exposure, but the BC-induced changes could be a key contributing factor in how air pollutants cause increased lower respiratory tract infectious disease.

Here we report investigation of not only the impact of BC on nasopharyngeal colonisation and invasion by the community-acquired, methicillin resistant *S. aureus* (CA-MRSA) strain USA300 LAC but also the genetic mechanisms involved. We show that, relative to *S. aureus* alone, simultaneous inoculation of BC and *S. aureus* into the nasopharynx increases *S. aureus* numbers in the nasopharynx, lungs, and blood of mice, and increases *S. aureus* adhesion to human respiratory epithelial cells. *S. aureus* grown in BC prior to inoculation also showed increased murine respiratory tract colonisation and invasion *in vivo* as well as adhesion and invasion of human epithelial cells *in vitro* supporting the hypothesis that BC acts as a hitherto unconsidered signal that has a direct effect on *S. aureus* behaviour. Global transcriptional analysis showed that BC does indeed have a widespread effect on *S. aureus* transcriptional responses, altering the regulation of the major virulence gene regulators Sae and Agr and causing increased expression of genes for toxins, proteases, and immune evasion factors.

## Results

### Exposure of S. aureus to BC prior to inoculation increases bacterial numbers in the respiratory tract

Previous studies have shown that bacterial dissemination from the nasopharynx to murine or rat lungs is induced when the animals are exposed to different forms of PM before bacterial inoculation or when bacteria and PM are simultaneously inoculated (Hussey et al., 2017; Shears et al., 2020; Yadav et al., 2020; Zhao et al., 2014). However, the direct effect of PM on bacterial behaviour during colonisation was not fully established because the presence of significant levels of PM within the host could potentiate colonisation by several mechanisms including damaging host tissue, by acting as a vehicle to support bacterial dissemination through the respiratory tract, or by supporting bacterial growth.

Our previous work showed that BC increased colonisation and invasion by *S. pneumoniae*, however it was not established whether this was due to direct effects on the bacterium and/or to effects on the host. Biologically relevant concentrations of BC are used that are at the lower end of the range of total amounts (96 μg – 378 μg per day) of air pollution particulate matter reported to be inhaled and deposited in the human respiratory tract each day (Chalvatzaki et al., 2018). To establish if BC directly affects the *in vivo* behaviour of *S. aureus*, mice were intranasally inoculated with the bacterium grown in the presence of BC but with the BC particles removed by dilution prior to inoculation. Mice were also inoculated with *S. aureus* LAC alone and LAC simultaneously inoculated with BC.

*S. aureus* pre-grown in BC prior to inoculation (gBC) significantly increased staphylococci in the nasopharynx (Fig. 1A) and lungs (Fig. 1B) at day 7 post-infection period compared to the control without BC (p<0.05). When *S. aureus* were co-inoculated together with BC (LAC+BC) a significant increase in *S. aureus* in the nasopharynx and lungs by 7 days post-infection was also observed (Fig. 1A and B both p<0.01 compared to the control). In contrast, only after co-inoculation of BC and *S. aureus* were there more staphylococci in the blood compared to the BC control (Fig. 1C; p<0.05). None of the mice had visible signs of disease and all survived throughout the experiment. Together these data indicate that the increase in infection of the nasopharynx and lungs by *S. aureus* is caused by the BC acting directly on the bacterium rather than via the host, but the presence of BC is important for bacterial invasion to the blood.

**Figure 1.**
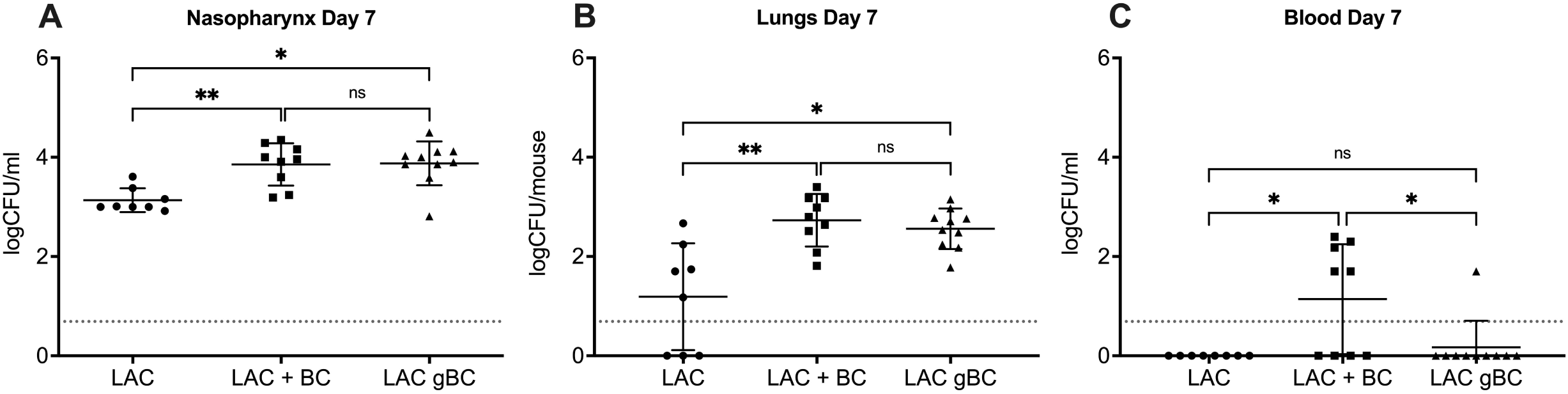
Exposure of *S. aureus* to BC results in increased respiratory tract colonisation in mice. Female CD1 mice were intranasally inoculated with 15μl containing 10^7^ *S. aureus* LAC (-BC), 10^7^ LAC with 105 μg BC (+BC) or 10^7^ CFU LAC pre-grown in the presence of 100 μg/ml BC (gBC). After 7 days, bacteria were recovered from the nasopharynx (A), lungs (B) and blood (C) and plated to determine the bacterial CFUs. No mice showed any clinical signs of infection. Data are presented as logCFU/ml for nasopharyngeal washes and blood, and logCFU/mouse for lungs. The dotted line marks the limit of detection. Data were analysed using a Kruskal-Wallis Test with Dunn’s multiple comparison (* = p<0.05, **= p<0.01).

### S. aureus pre-grown in BC show increased adhesion and invasion of epithelial cells in vitro

Because *S. aureus* can invade non-professional phagocytes thereby avoiding aspects of the immune system (Garzoni & Kelley, 2008), we determined whether BC alters adherence or invasion of *S. aureus* to human respiratory epithelial cells. The type II-like bronchial epithelial cells, A549, were exposed to *S. aureus* LAC grown in the same conditions as the murine colonisation.

There was a significant increase in numbers of *S. aureus* adhering to (Fig. 2A p<0.0001) and invading (Fig. 2B p<0.01) A549 cells when grown with BC (gBC) prior to inoculation compared to bacteria grown in the absence of BC (-BC), an observation consistent with a direct effect of BC on the bacteria. There was also increased adhesion of *S. aureus* to A549 cells when simultaneously inoculated with BC (+BC) (Fig. 2A p<0.01) but no significant change in invasion. Importantly inoculation with BC alone did not affect A549 cell viability (Fig. 2C). Incubation of A549 with *S. aureus* LAC caused 18% cytotoxicity but there was no significant increase in cytotoxicity on A549 cells during co-inoculation with BC or when *S. aureus* were pre-grown in BC (Fig. 2C). Together these data demonstrate that increased *S. aureus* epithelial adhesion and invasion is caused by direct interaction of BC with the bacteria and does not involve gross changes to the epithelial cells.

**Figure 2.**
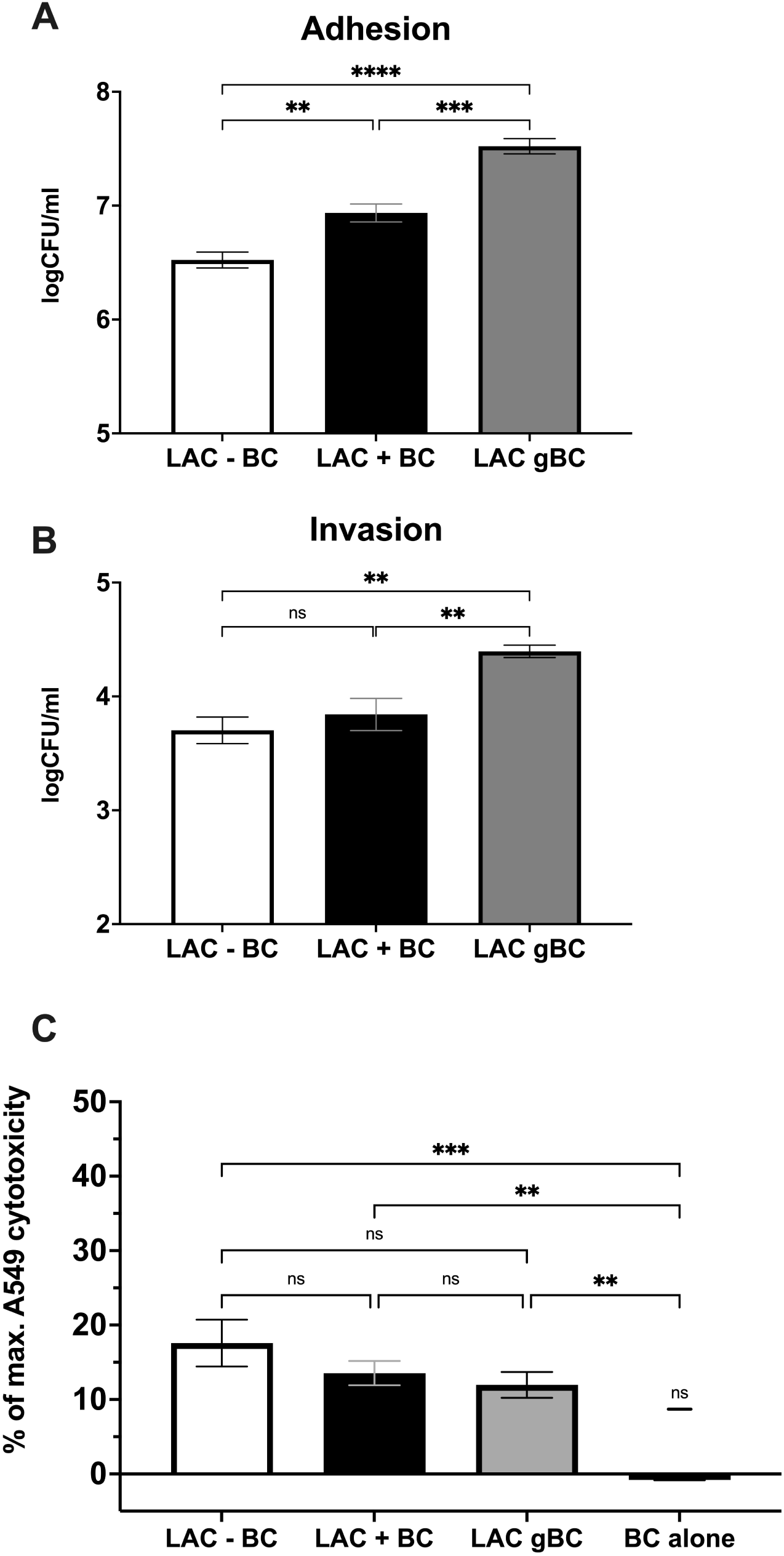
Exposure to BC results in increased adhesion, invasion and persistence within human epithelial cells. *S. aureus* LAC adhesion (A), and invasion (B) of human lung epithelial A549 cells was measured using a gentamicin protection assay. Monolayers of 1×10^5^ A549 cells in 24-well plates were infected at a MOI of 100. Data are presented as logCFU/ml and error bars represent 1SEM of at least 5 biological repeats. Significance was determined by one-way ANOVA with Dunnett’s multiple comparison test (* = p<0.05, **= p<0.01, ***=P<0.001, ****=p<0.0001). (C) Cytotoxicity was measured through lactate dehydrogenase (LDH) release from A549 cells after 2 hours exposure to *S. aureus* and/or BC. % cytotoxicity is calculated relative to spontaneous cell death (0%) and maximum cell death (100% cell lysis). Significance was determined by One-way ANOVA with Tukey’s multiple comparison test (* = p<0.05, **= p<0.01, ***=P<0.001, ****=p<0.0001).

### BC induces expression of S. aureus genes for toxins and proteases and the SOS response

Having shown that BC alters the phenotype of *S. aureus*, we used transcriptome sequencing (RNAseq) to investigate the pattern of gene expression induced using the same BC exposure growth conditions as the colonisation experiments. RNAseq analysis identified 52 staphylococcal genes that showed a significant increase in expression (Table 1 and S2) and 63 genes that showed a significant decrease in expression (Table 2 and S3) after exposure to BC.

**Table 1.** Genes significantly upregulated at a Fold change > 2 in response to BC. Adjusted pValues for all genes are < 0.001.

| Locus Tag<br>(SAUSA300) | Gene<br>name | Product Description | Fold<br>Change |
| --- | --- | --- | --- |
| RS11190 | *kdpA* | potassium-transporting ATPase subunit A | 6.30 |
| RS09620 | *splA* | serine protease SplA | 4.88 |
| RS10530 | *chp* | chemotaxis-inhibiting protein CHIPS | 4.75 |
| RS09610 | *splC* | serine protease SplC | 4.40 |
| RS09615 | *splB* | serine protease SplB | 4.04 |
| RS09595 | *splF* | serine protease SplF | 3.99 |
| RS05720 | *hla* | alpha-hemolysin | 3.81 |
| RS09605 | *splD* | serine protease SplD | 3.75 |
| RS10525 | *scn* | complement inhibitor SCIN-A | 3.60 |
| RS15090 | - | phenol-soluble modulins PSM-alpha-3 | 3.53 |
| RS15730 | - | phenol-soluble modulins PSM-alpha-4 | 3.50 |
| RS05795 | *psm_2* | beta-class phenol-soluble modulins | 3.45 |
| RS15735 | - | phenol-soluble modulins PSM-alpha-2 | 3.44 |
| RS00185 | - | DUF1643 domain-containing protein | 3.29 |
| RS15740 | - | phenol-soluble modulins PSM-alpha-1 | 3.07 |
| RS06445 | *glpD* | glycerol-3-phosphate dehydrogenase/oxidase | 2.96 |
| RS01705 | *gehB* | YSIRK domain-containing triacylglycerol lipase Lip2/Geh | 2.90 |
| RS13080 | *hlgB* | bi-component gamma-hemolysin HlgAB/HlgCB subunit B | 2.87 |
| RS06715 |  | hypothetical protein | 2.86 |
| RS09600 | *splE* | serine protease SplE | 2.83 |
| RS06840 | *mucB* | DNA repair protein MucB | 2.78 |
| RS00515 | *plc* | phosphatidylinositol-specific phospholipase C | 2.68 |
| RS05790 | *psm_1* | beta-class phenol-soluble modulins | 2.68 |
| RS02340 | *metQ2* | dipeptide ABC transporter glycylmethionine-binding lipoprotein | 2.63 |
| RS01835 | *fepC* | iron permease FTR1 family protein | 2.61 |
| RS06370 | *recA* | DNA recombination/repair protein RecA | 2.61 |
| RS04005 | *uvrA* | excinuclease ABC subunit UvrA | 2.61 |
| RS04395 | *ear* | DUF4888 domain-containing protein | 2.57 |
| RS02335 | *metP2* | ABC transporter permease | 2.53 |
| RS14170 | *nrdG* | anaerobic ribonucleoside-triphosphate reductase activating protein | 2.46 |
| RS04985 | *comK1* | competence protein ComK | 2.42 |
| RS13075 | *hlgC* | bi-component gamma-hemolysin HlgCB subunit C | 2.36 |
| RS10660 | - | HNH endonuclease | 2.34 |
| RS10495 |  | MAP domain-containing protein | 2.30 |
| RS10845 | *lukG* | bi-component leukocidin LukGH subunit G | 2.29 |
| RS01825 | *fepA* | EfeM/EfeO family lipoprotein | 2.27 |
| RS02330 | *metN2* | methionine ABC transporter ATP-binding protein | 2.15 |
| RS10505 | *hly-1* | sphingomyelin phosphodiesterase | 2.14 |
| RS04000 | *uvrB* | excinuclease ABC subunit UvrB | 2.13 |
| RS14555 | - | S-adenosyl-L-methionine hydroxide adenosyltransferase family protein | 2.13 |
| RS06710 | *lexA* | transcriptional repressor LexA | 2.11 |
| RS13060 | *sbi* | immunoglobulin-binding protein Sbi | 2.11 |
| RS10420 | - | YolD-like family protein | 2.10 |
| RS07540 |  | Panton-Valentine bi-component leukocidin subunit F | 2.09 |
| RS08875 | *infC* | translation initiation factor IF-3 | 2.09 |
| RS04990 | - | IDEAL domain-containing protein | 2.07 |
| RS03850 | *nrdF* | class 1b ribonucleoside-diphosphate reductase subunit beta | 2.05 |
| RS03840 | *nrdI* | class 1b ribonucleoside-diphosphate reductase assembly flavoprotein NrdI | 2.05 |
| RS11200 | *kdpD* | sensor histidine kinase KdpD | 2.04 |
| RS08675 | *recJ* | single-stranded-DNA-specific exonuclease RecJ | 2.04 |
| RS06750 | *sbcC* | SMC family ATPase | 2.04 |
| RS10850 | *lukH* | bi-component leukocidin LukGH subunit H | 2.03 |

**Table 2.**
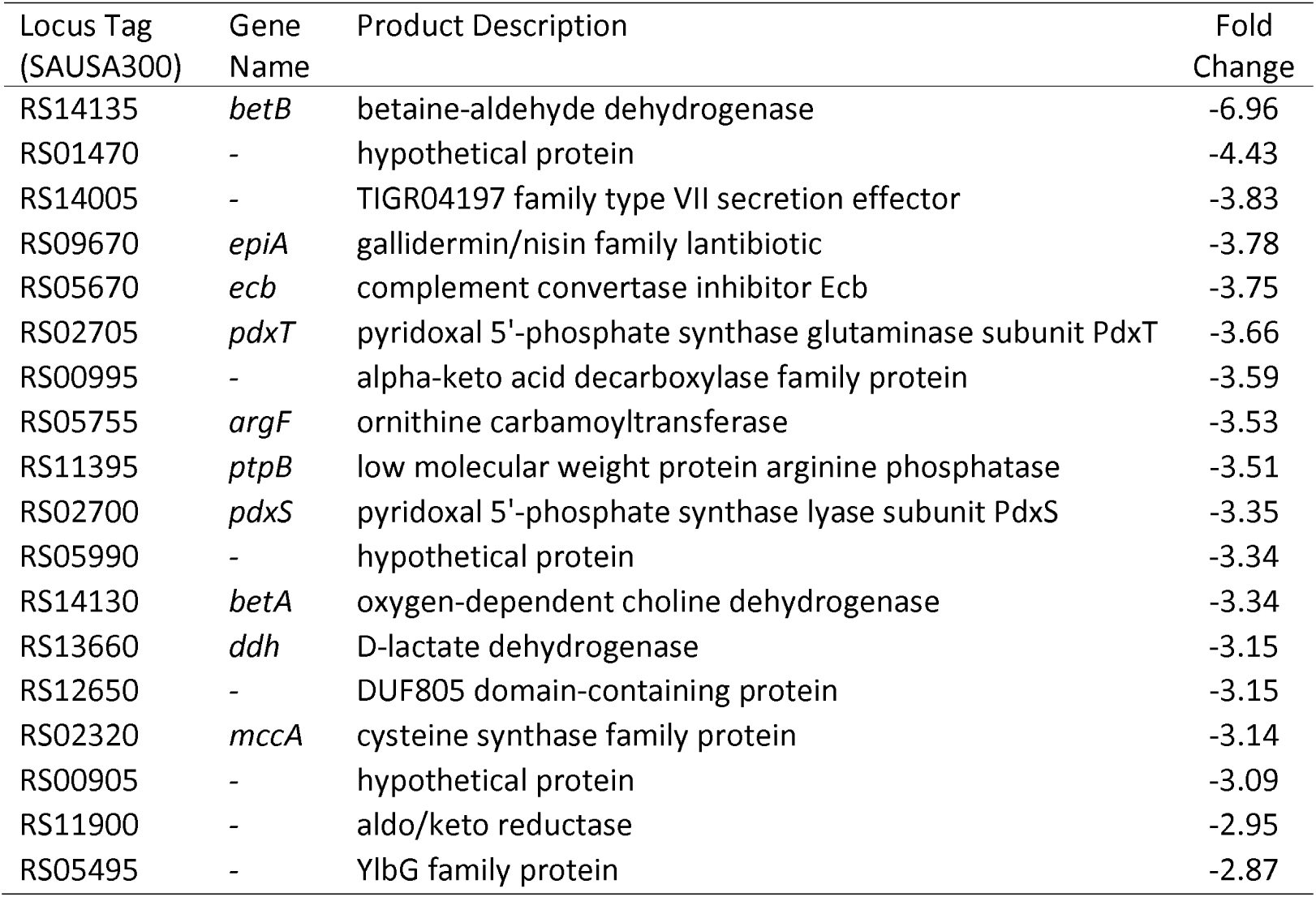
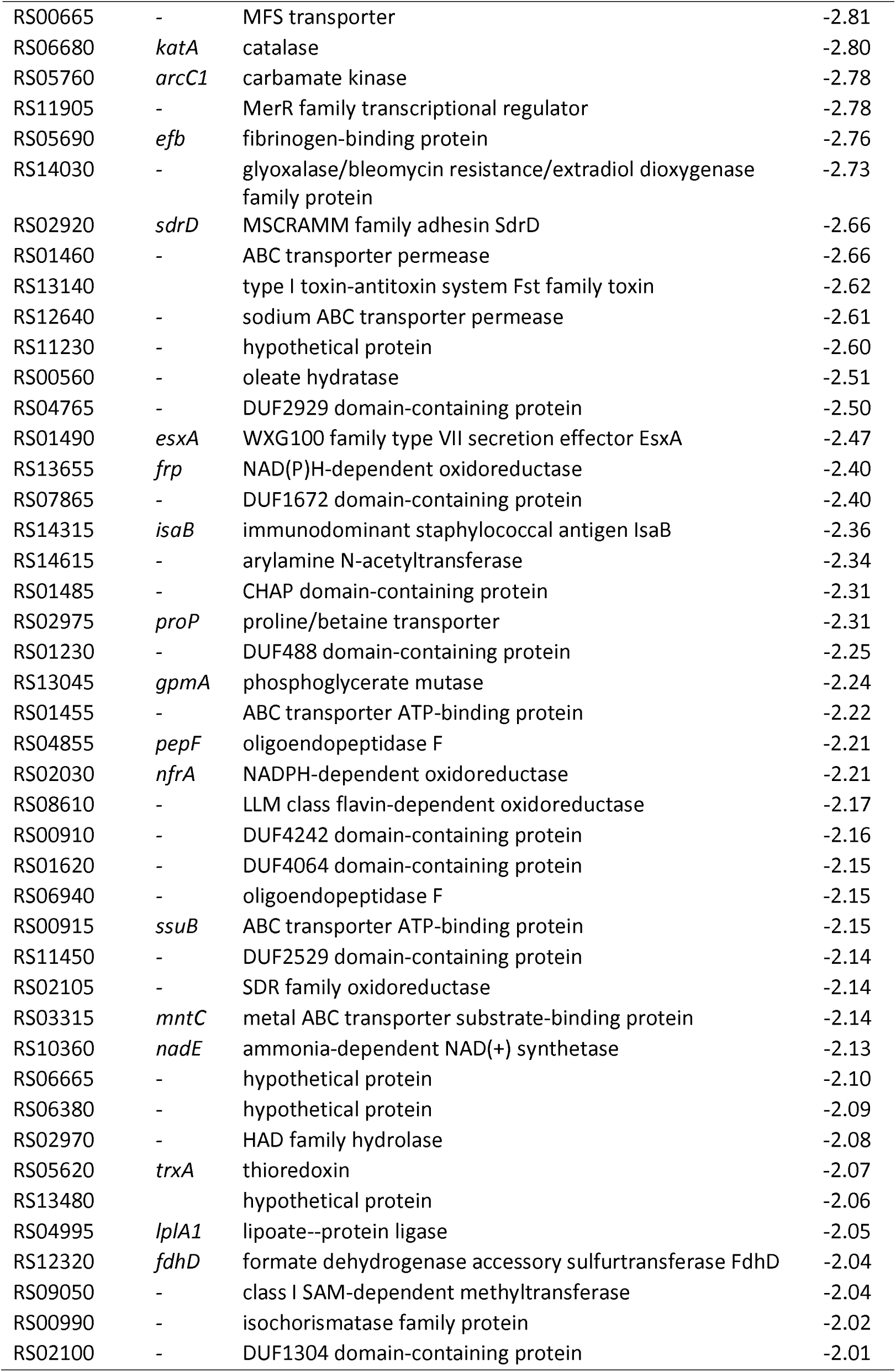
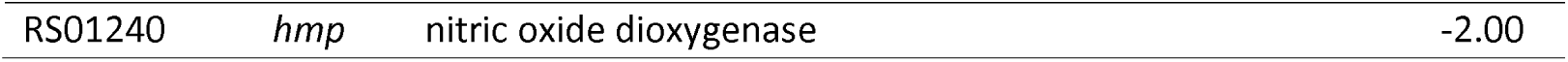
Genes significantly downregulated at a Fold Change < -2 in response to BC. Adjusted pValues for all genes are < 0.001.

| Locus Tag<br>(SAUSA300) | Gene<br>Name | Product Description | Fold<br>Change |
| --- | --- | --- | --- |
| RS14135 | *betB* | betaine-aldehyde dehydrogenase | -6.96 |
| RS01470 | - | hypothetical protein | -4.43 |
| RS14005 | - | TIGR04197 family type VII secretion effector | -3.83 |
| RS09670 | *epiA* | gallidermin/nisin family lantibiotic | -3.78 |
| RS05670 | *ecb* | complement convertase inhibitor Ecb | -3.75 |
| RS02705 | *pdxT* | pyridoxal 5'-phosphate synthase glutaminase subunit PdxT | -3.66 |
| RS00995 | - | alpha-keto acid decarboxylase family protein | -3.59 |
| RS05755 | *argF* | ornithine carbamoyltransferase | -3.53 |
| RS11395 | *ptpB* | low molecular weight protein arginine phosphatase | -3.51 |
| RS02700 | *pdxS* | pyridoxal 5'-phosphate synthase lyase subunit PdxS | -3.35 |
| RS05990 | - | hypothetical protein | -3.34 |
| RS14130 | *betA* | oxygen-dependent choline dehydrogenase | -3.34 |
| RS13660 | *ddh* | D-lactate dehydrogenase | -3.15 |
| RS12650 | - | DUF805 domain-containing protein | -3.15 |
| RS02320 | *mccA* | cysteine synthase family protein | -3.14 |
| RS00905 | - | hypothetical protein | -3.09 |
| RS11900 | - | aldo/keto reductase | -2.95 |
| RS05495 | - | YlbG family protein | -2.87 |
| RS00665 | - | MFS transporter | -2.81 |
| RS06680 | *katA* | catalase | -2.80 |
| RS05760 | *arcC1* | carbamate kinase | -2.78 |
| RS11905 | - | MerR family transcriptional regulator | -2.78 |
| RS05690 | *efb* | fibrinogen-binding protein | -2.76 |
| RS14030 | - | glyoxalase/bleomycin resistance/extradiol dioxygenase family protein | -2.73 |
| RS02920 | *sdrD* | MSCRAMM family adhesin SdrD | -2.66 |
| RS01460 | - | ABC transporter permease | -2.66 |
| RS13140 |  | type I toxin-antitoxin system Fst family toxin | -2.62 |
| RS12640 | - | sodium ABC transporter permease | -2.61 |
| RS11230 | - | hypothetical protein | -2.60 |
| RS00560 | - | oleate hydratase | -2.51 |
| RS04765 | - | DUF2929 domain-containing protein | -2.50 |
| RS01490 | *esxA* | WXG100 family type VII secretion effector EsxA | -2.47 |
| RS13655 | *frp* | NAD(P)H-dependent oxidoreductase | -2.40 |
| RS07865 | - | DUF1672 domain-containing protein | -2.40 |
| RS14315 | *isaB* | immunodominant staphylococcal antigen IsaB | -2.36 |
| RS14615 | - | arylamine N-acetyltransferase | -2.34 |
| RS01485 | - | CHAP domain-containing protein | -2.31 |
| RS02975 | *proP* | proline/betaine transporter | -2.31 |
| RS01230 | - | DUF488 domain-containing protein | -2.25 |
| RS13045 | *gpmA* | phosphoglycerate mutase | -2.24 |
| RS01455 | - | ABC transporter ATP-binding protein | -2.22 |
| RS04855 | *pepF* | oligoendopeptidase F | -2.21 |
| RS02030 | *nfrA* | NADPH-dependent oxidoreductase | -2.21 |
| RS08610 | - | LLM class flavin-dependent oxidoreductase | -2.17 |
| RS00910 | - | DUF4242 domain-containing protein | -2.16 |
| RS01620 | - | DUF4064 domain-containing protein | -2.15 |
| RS06940 | - | oligoendopeptidase F | -2.15 |
| RS00915 | *ssuB* | ABC transporter ATP-binding protein | -2.15 |
| RS11450 | - | DUF2529 domain-containing protein | -2.14 |
| RS02105 | - | SDR family oxidoreductase | -2.14 |
| RS03315 | *mntC* | metal ABC transporter substrate-binding protein | -2.14 |
| RS10360 | *nadE* | ammonia-dependent NAD(+) synthetase | -2.13 |
| RS06665 | - | hypothetical protein | -2.10 |
| RS06380 | - | hypothetical protein | -2.09 |
| RS02970 | - | HAD family hydrolase | -2.08 |
| RS05620 | *trxA* | thioredoxin | -2.07 |
| RS13480 |  | hypothetical protein | -2.06 |
| RS04995 | *lplA1* | lipoate--protein ligase | -2.05 |
| RS12320 | *fdhD* | formate dehydrogenase accessory sulfurtransferase FdhD | -2.04 |
| RS09050 | - | class I SAM-dependent methyltransferase | -2.04 |
| RS00990 | - | isochorismatase family protein | -2.02 |
| RS02100 | - | DUF1304 domain-containing protein | -2.01 |
| RS01240 | *hmp* | nitric oxide dioxygenase | -2.00 |

Gene Ontology (GO) Enrichment Analysis was used to determine which GO groupings of biological processes (BP), molecular functions (MF) and cellular components (CC) were statistically over-represented in the presence of BC compared with the absence of BC. The genes upregulated in response to BC showed significant over-representation of 19 BP and 6 MF genes (Fig. 3). In the BP category the top 4 terms (cell killing GO:0001906, cytolysis in other organisms GO:0051715, killing of cells of other organisms GO:0031640 and haemolysis in another organism GO:0044179) are all involved in cell killing. This is mirrored in the MF, in that the most overrepresented term is toxin activity (GO:0090729). Other BPs that show over-representation include those involved in the response to environmental changes and DNA damage and repair. There was no significant over-representation of any biological processes or molecular functions in the negatively regulated genes and no cellular components in either growth condition.

**Figure 3.**
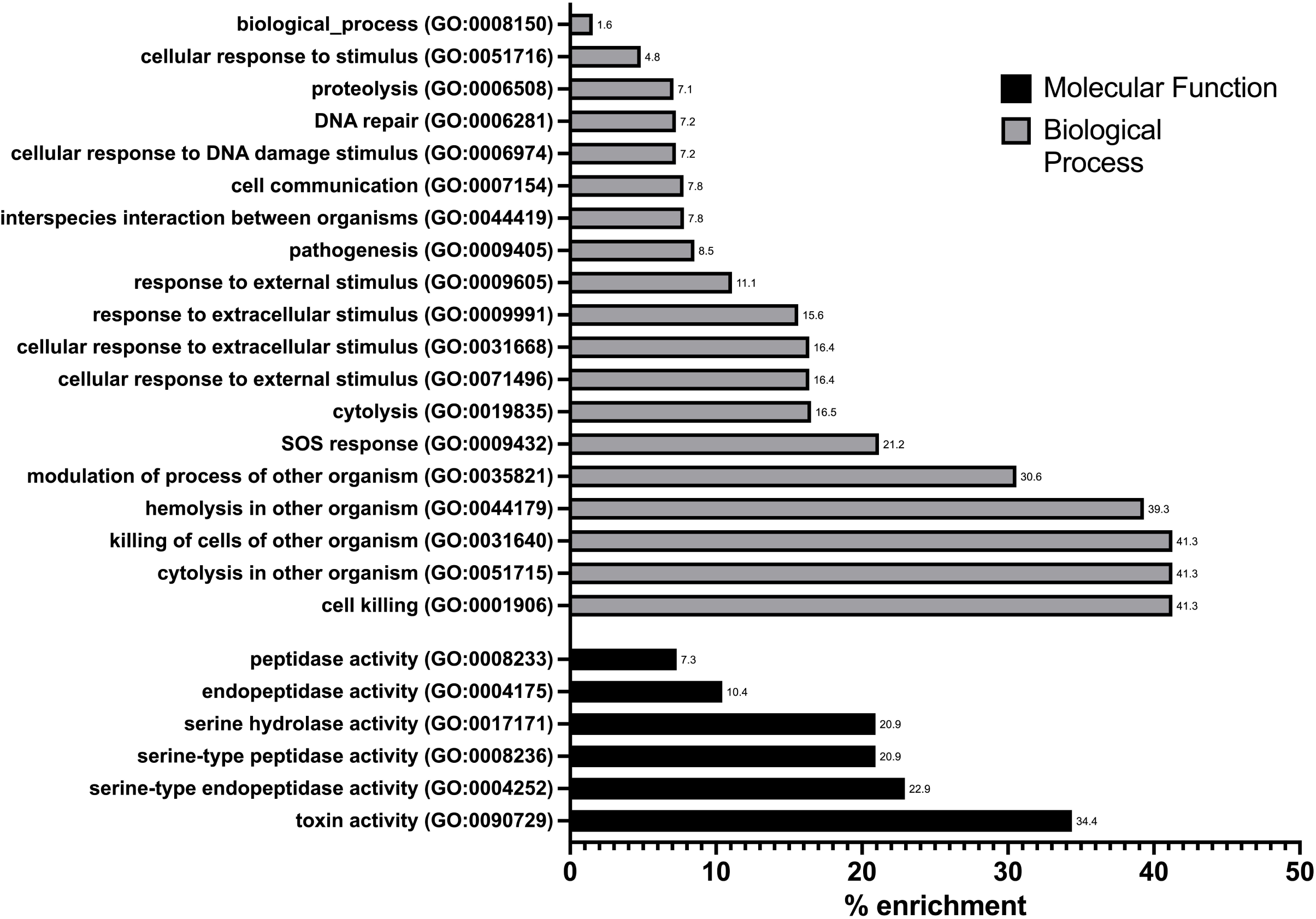
BC exposure results in increased expression of genes involved in toxin production, proteases and DNA replication and repair. Gene Ontology (GO) enrichment analysis (A) of 52 genes upregulated in response to BC. The chart shows the over-representation of each GO term within the dataset as a % enrichment.

The differentially expressed genes were also grouped based on their main function, using the associated TIGRFAM number which automatically groups proteins, based on sequence homology, into functional families and provides most-likely functions for hypothetical and unannotated genes (Haft et al., 2013; Haft et al., 2018)(Tables S2 and S3, Fig. 4). Of the 52 genes upregulated in response to BC, 16 (30.1%) are involved in toxin production, immune evasion, or pathogenesis, 11 genes (21.1%) are involved in DNA metabolism, replication, recombination and repair, and 8 genes (15%) play a role in protein synthesis, degradation and repair (Fig. 4A). The remaining genes mainly play roles in cellular processes, cell envelope and signalling, transport and binding, and 6 genes (11%) are currently undefined. Of the 63 genes downregulated in response to BC, the largest represented groups contain 10 genes (15.9%) involved in central metabolism and 8 genes (12.7%) involved in energy metabolism, with 22 undefined genes (35%) (Fig. 4B).

**Figure 4.**
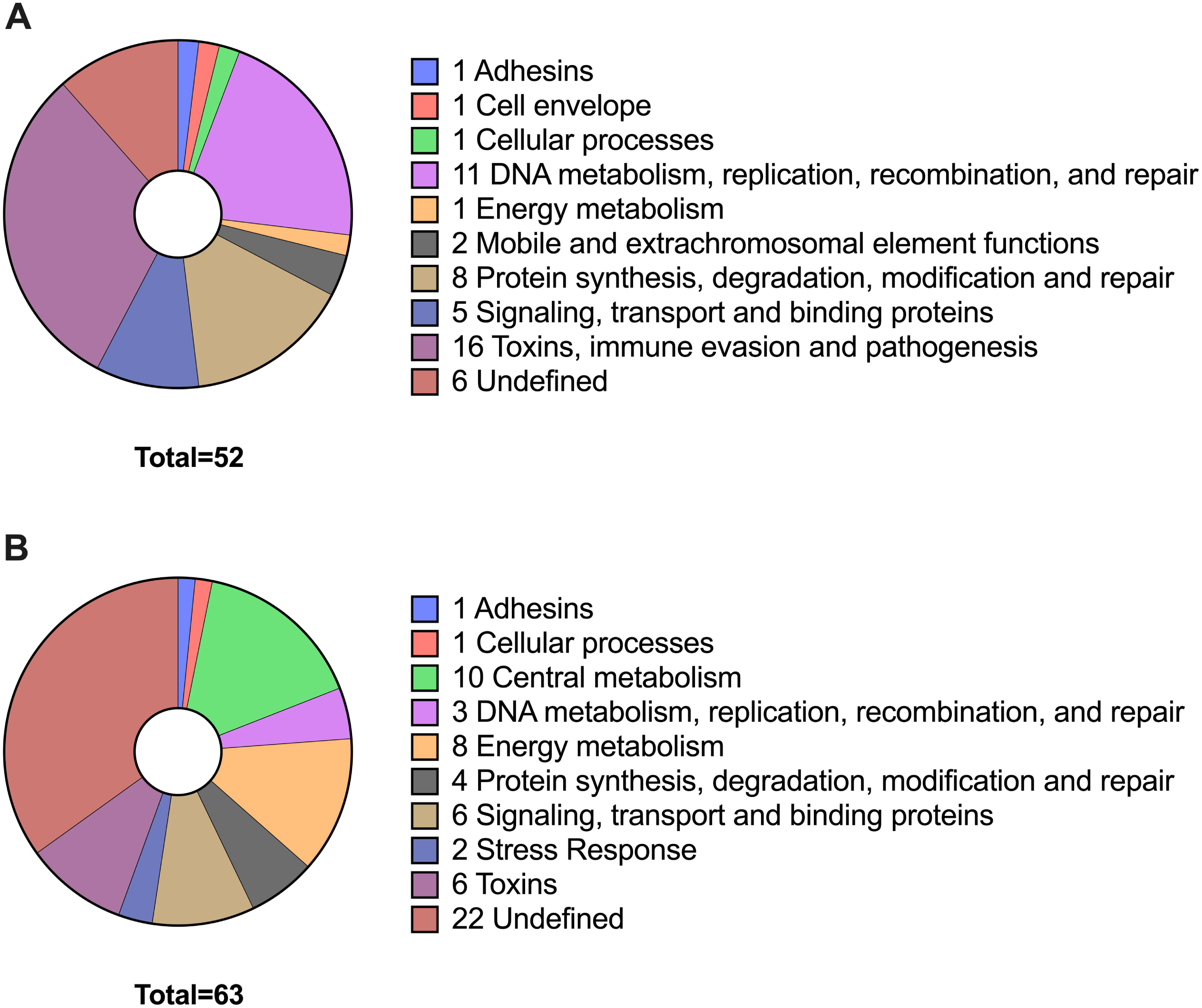
BC exposure results in differential regulation in *S. aureus*. Pie chart organising (A) the 52 up-regulated genes and (B) the 63 down-regulated genes, by their main TIGRFAM function.

GO analysis showed that genes associated with pathogenesis are induced by BC. We observed 2 to 6-fold induction of the serine protease genes (*splA, B, C, D, E, F*), and genes for toxins and immune evasion (*hla, hlb-1, hlgBC, gehB, scn, chp, plc, psm alpha-1, 2, 3, 4, psm beta-1, 2, sbi, lukF-PV, lukG, H)*. It is notable that all these genes are regulated by either the Agr quorum sensing system (Bronesky et al., 2016; Kavanaugh & Horswill, 2016; Le & Otto, 2015) or the SaeRS two component regulatory system (Liu, Q. et al., 2016; Voyich et al., 2009) or both (Table 1).

Several genes involved in DNA repair show a 2 to 2.6-fold induction in response to BC exposure. These include the genes for the UvrAB nucleotide excision repair endonuclease, the UmuC error-prone polymerase V and ribonucleotide reductase, HNH endonuclease, Yol-D family protein and single-stranded-DNA-specific exonuclease RecJ that form an integral part of the SOS response (Podlesek & Žgur Bertok, 2020).

The other genes induced by BC are for glycerol-3-phosphate dehydrogenase/oxidase (*glpD*), the FepABC haem utilisation system (*fepAC*) (Turlin et al., 2013), the dipeptide methionine transporter (*metNPQ2*) (Wade et al., 2004) and the potassium transporter and regulator (*kdpA, kdpD*) (Xue et al., 2011). These genes are regulated by CcpA, the iron repressor protein Fur, the cysteine metabolism regulator CymR and KdpDE two component regulator respectively (Fuchs et al., 2018; Nagarajan & Elasri, 2007).

### BC represses expression of genes for stress responses and metabolism in S. aureus

GO analysis did not show any significant over-representation of down-regulated genes, unlike the up-regulated genes. Several genes that are repressed in BC are typically induced in response to different stresses, including those involved in oxidative stress (*katA, trxA*), osmotic stress (glycine betaine synthesis *betAB*, proline/betaine transporter *proP*), sulphur metabolism (*cysM, proP, ssuB*) and nitrosative stress (*hmp, ldh*).

Exposure to BC also results in repression of genes for some adhesins (*sdrD, efb)* and an immune evasion factor (*ecb*). Interestingly, although the stress response (*betAB, ldh, proP*) and metabolic genes (*argF, ptpB, pepF, nadE*) that are repressed by Agr are also repressed by BC, the majority of the adhesin genes normally repressed by Agr showed no change in expression (e.g. *fnbA, fnbB, emp, spa;* refs), suggesting that the Agr regulon is only partially affected by BC. It is also noteworthy that BC exposure partially repressed the regulons of other global regulators (e.g. CymR (*cysM*), SigB (*proP*), GraRS (*ldh, entB*) PdxR (*pdxS, pdxT*), RexAB (*frp, ldh*), PerR (*katA, trxA*) (Fuchs et al., 2018; Nagarajan & Elasri, 2007)) showing that BC acts as a signal that induces a newly described pattern of *S. aureus* global gene expression.

To establish whether BC induces gene expression at lower BC concentrations, qRT-PCR was used to determine *S. aureus* gene expression in response to BC at 5, 50 and 100 μg ml^-1^. The expression of the virulence genes *chp, splF, betB, ecb*, and *epiA* all showed a clear concentration-dependent effect in response to BC (Fig. 5) showing that BC can induce *S. aureus* gene expression at low concentrations.

**Figure 5.**
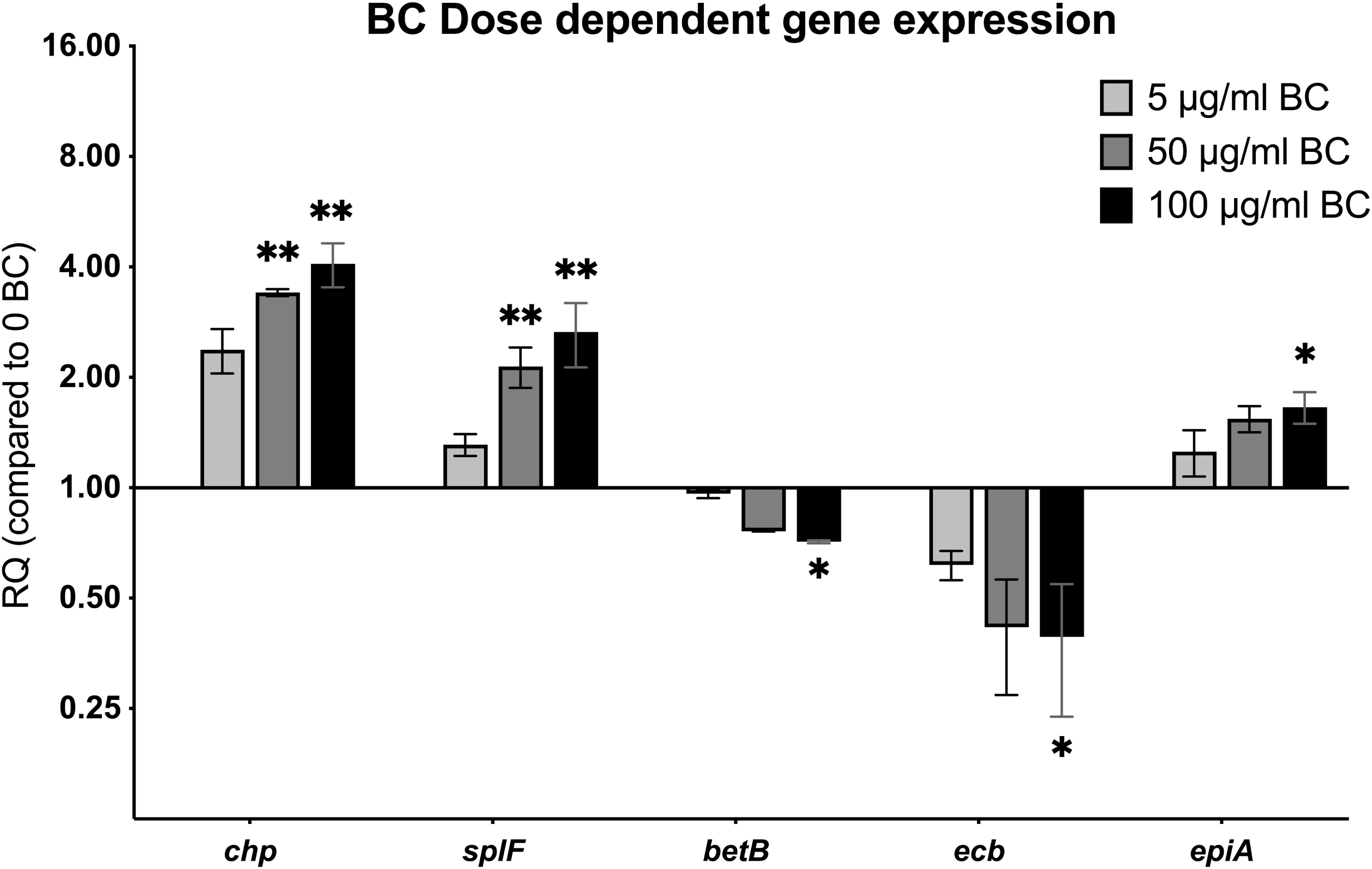
Gene expression changes in response to BC are dose dependent. Relative fold change in *S. aureus* USA300 gene expression grown in the presence of 5, 50 and 100 μg/ml BC. RQ is the fold change in expression relative to – BC. Significance of each concentration compared to 0 BC was determined by Kruskal Wallis Test with Dunn’s multiple comparison test (* = p<0.05, **= p<0.01).

### BC induced transcriptional changes require functional Agr and Sae regulators

The transcriptional analysis showed that BC alters the expression of only subsets of the Agr and Sae regulons. This conclusion from the RNAseq analysis was tested using qRT-PCR. Of the Agr and Sae responsive genes investigated, there were significant changes in expression of the Agr (*hla;* Fig. 6 p<0.05) and Sae (*chp;* Fig. 6 p<0.01) regulated genes for toxin and immune evasion factors, in agreement with the RNAseq data, but none of the other Agr- and Sae-regulated adhesins (*clfA, fnbA, fnbB, emp*) showed significant change in response to BC (p>0.05), also in agreement with the RNA seq analysis (Fig. 6). Furthermore, PerR-regulated oxidative stress genes (Horsburgh et al., 2001) also showed differential regulation in response to BC, with decreased transcription of *katA* (Fig. 6 p<0.05) but no change in that of the *sodA* gene. Overall, these data confirm the RNAseq analysis and show that BC induces a hitherto unseen pattern of gene expression.

**Figure 6.**
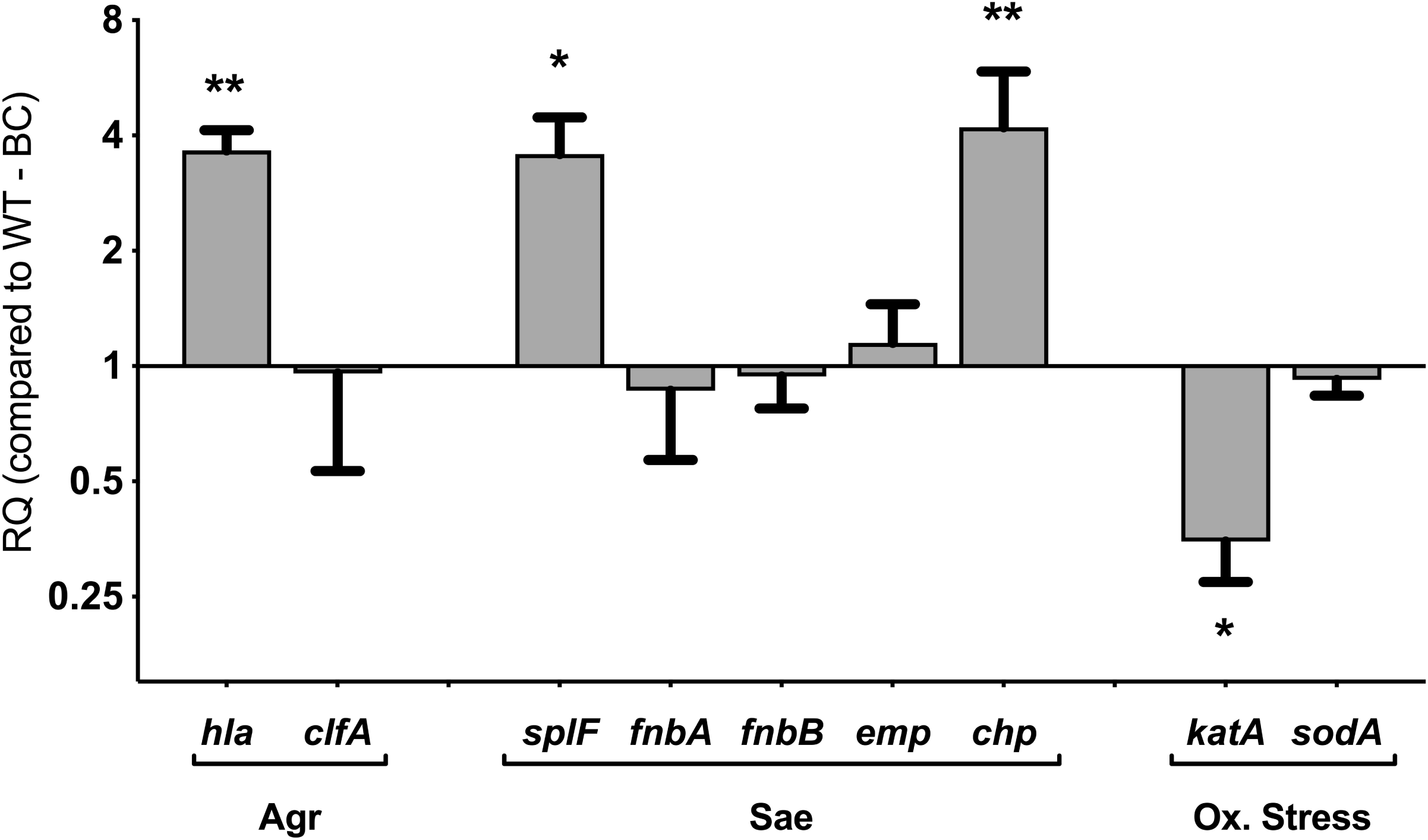
BC induces unexpected patterns of gene expression in key virulence and stress regulons. Relative fold change in *S. aureus* USA300 gene expression grown in the presence of 100 μg/ml BC. Effector genes are grouped based on their primary regulator (Agr, Sae or Oxidative stress). RQ is the fold change in expression relative to – BC. Significance of each concentration compared to 0 BC was determined by Kruskal Wallis Test with Dunn’s multiple comparison test (* = p<0.05, **= p<0.01).

To investigate the role of the Agr and SaeRS regulators in the bacterial response to BC, *S. aureus* LAC *agrB* and *saeS* transposon insertion mutants were constructed as described in the methods. With these *S. aureus* LAC mutants, the transcription of the Agr-regulated *psmβ*, the Sae-regulated *chp* and the dual Agr- and Sae-regulated *lukS-PV* and *splF* genes in response to BC was investigated. The expression of *kdpD* was also investigated because *kdpD* is indirectly induced by Agr through repression of the repressor Rot, but it is not regulated by Sae (Xue et al., 2011). In the absence of BC, transcriptional analysis confirmed previous studies of Agr and Sae regulation of these genes (Cheung et al., 2011; Liu et al., 2016). The expression of *lukS* and *splF* show 2-fold decrease in expression in both the *agr* and *sae* mutants, the *psmβ* gene was not expressed in the *agr* mutant, while *chp* was not expressed in the *sae* mutant, and *kdpD* showed no major change in expression in either the *agr* or *sae* mutants (Fig. 7).

**Figure 7.**
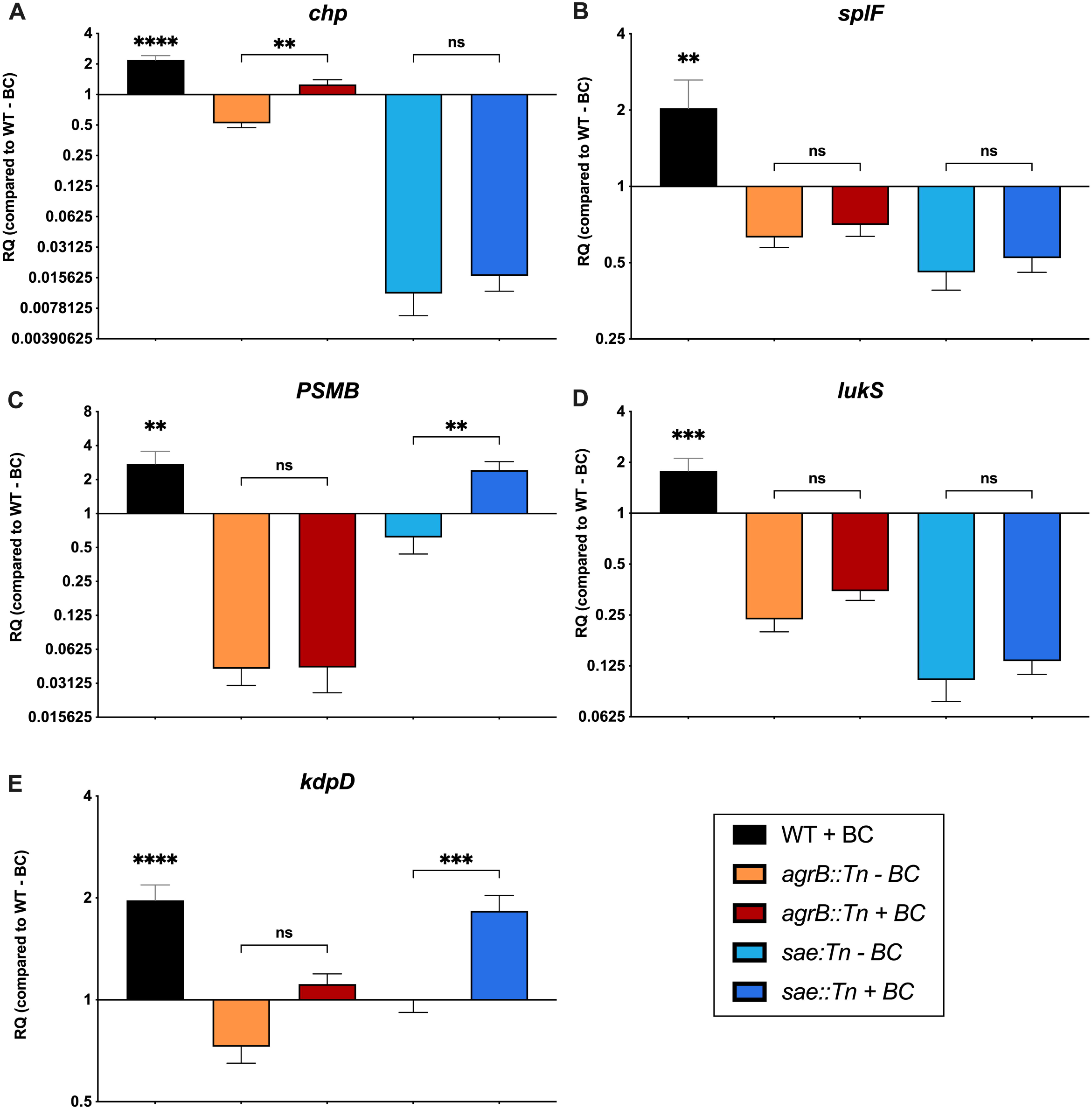
The transcriptional regulators *agr* and *sae* are involved in the BC regulation of some but not all BC induced genes. Transcriptional response of *S. aureus* genes in response to BC in USA300 LAC WT, *agrB*::Tn and *sae*::Tn mutant strains. RQ is the fold change in expression in each strain relative to the WT – BC. Significance was determined by One-way ANOVA with Tukey’s multiple comparison test (* = p<0.05, **= p<0.01, ***=P<0.001, ****=p<0.0001).

Interestingly, in the presence of BC, significant induction of *chp* gene transcription remained in the *agr* mutant (Fig. 7A p<0.01) and *psmβ* and *kdpD* were still significantly induced by BC in the *sae* mutant (Fig. 7C p<0.01; Fig. 7E P<0.001). In contrast, there was no significant BC induction of the *lukS, psmβ* or *splF* gene expression in the *agr* mutant or *chp, splF* and *lukS* in the *sae* mutant. It is noteworthy that these data show that either Agr or Sae are required for BC induction of *chp, kdpD* and *psmβ* and that the BC response is facilitated by both genes either together or separately.

### BC induction of S. aureus epithelial cell invasion is via a sae-independent mechanism

Our data show that BC increases *S. aureus* adhesion and invasion to human epithelial cells. The ability of *S. aureus* to adhere to and invade non-professional phagocytes has been reported to be dependent on Sae induction of the adhesin genes *fnbB, fnbA, eap* and *atl* (Hirschhausen et al., 2010; Liang et al., 2006). To investigate the role of Sae and Agr in the BC-mediated increase in *S. aureus* adhesion and invasion, A549 were exposed to wild type *S. aureus* LAC, and *agrB* and *sae* mutants pre-grown in the presence and absence of BC.

As shown in Fig. 8, although BC significantly increases *S. aureus* LAC adhesion to A549 cells (Fig. 8A p<0.05), neither the *agrB* nor *sae* mutants showed significant changes in adhesion compared to the wild type in the presence or absence of BC (Fig. 8A), although the *agr* mutant shows a small decrease in the response compared to LAC and the *sae* mutant. In the absence of BC, the *agr* mutant showed a significant increase in invasion (Fig. 8B p<0.05) whereas the *sae* mutant showed a significant decrease in invasion (Fig. 8B p<0.0001), confirming previous studies of the roles of Sae and Agr in *S. aureus* invasion of epithelial cells (Liang et al., 2006; Wesson et al., 1998). In the presence of BC, there was no significant change in *S. aureus* invasion in the *agr* mutant compared to the wild type (Fig. 8B). In contrast, there was a significant increase in BC induced *S. aureus* invasion in the *sae* mutant (Fig. 8B P<0.001) demonstrating that BC mediates staphylococci invasion via Sae and Agr-independent mechanisms.

**Figure 8.**
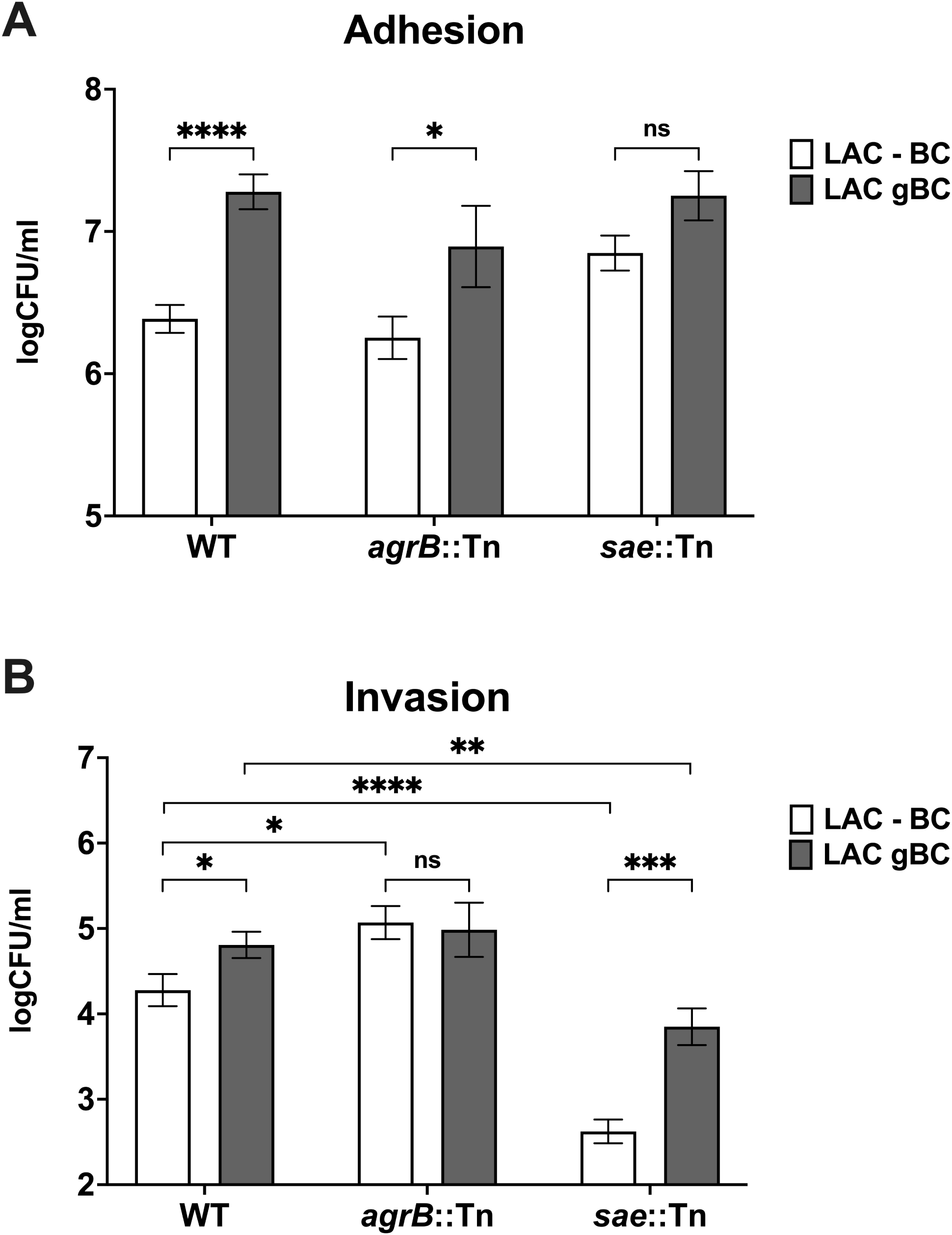
The effect of BC exposure on host cell interaction is involves Sae regulatory system. *S. aureus* LAC adhesion (A) and invasion (B) of human lung epithelial A549 cells by *S. aureus* LAC WT, *agrB*::Tn and *sae*::Tn mutants in response to BC was measured using a gentamycin protection assay. Cells were infected at a MOI of 100 on 1×10^5^ A549 monolayers in 24 well plates. Data is presented as logCFU/ml of recovered cells, and error bars represent 1SEM of at least 5 biological repeats. Significance was determined by 2-way ANOVA with Dunnett’s multiple comparison test (* = p<0.05, **= p<0.01, ***=P<0.001, ****=p<0.0001).

## Discussion

Black carbon (BC) is a major component of particulate matter (PM) in air pollution (Bell et al., 2007). Here we show that BC increases *S. aureus* colonisation of the murine respiratory tract and increases the bacterium’s adhesion to human respiratory epithelial cells and its invasion of these cells. Our data show that increased colonisation is due to the direct impact of BC on the bacteria and occurs in the absence of any detected BC mediated effects on the host. BC has a widespread effect on *S. aureus* global transcription causing increased expression of genes for toxins, proteases, and immune evasion factors critical for dissemination and colonisation. Together these data provide evidence for a new causative mechanism of the detrimental effects of air pollution in that air pollutants directly change bacterial gene expression altering their invasive capacity and their ability to colonise and disseminate within the respiratory tract.

There is growing epidemiological evidence that PM exposure increases the risk of infectious diseases that can be caused or exacerbated by *S. aureus;* for example, community acquired pneumonia, that is increasingly caused by CA-MRSA (Pivard et al., 2021), infective endocarditis (Hsieh et al., 2019), cystic fibrosis (Psoter et al., 2015; Psoter et al., 2017), chronic rhinosinusitis (Schwarzbach et al., 2020) and chronic skin diseases (Dijkhoff et al., 2020). Exposure to atmospheric PM has great potential to affect the activities of *S. aureus* because the bacterium persistently or transiently colonises the anterior nares and the skin (Pivard et al., 2021), where it will be exposed to PM.

In this study we show that simultaneous inoculation of *S. aureus* and biologically relevant concentrations of BC causes increased infection of murine lungs and increased nasopharyngeal colonisation. These new data agree with our previous observations with *S. pneumoniae* exposed to BC (Hussey et al., 2017) and confirms a wider phenomenon of the impact of BC on bacterial respiratory tract colonisation, that also been recently shown with other types of particulate pollutant (Liu et al., 2019; Shears et al., 2020; Woo et al., 2018; Yadav et al., 2020).

It is notable that we showed that pre-growth of *S. aureus* in BC prior to infection induces a significant increase in staphylococcal murine respiratory tract colonisation. The increased colonisation is maintained for at least 7-days without further administration of BC. To our knowledge, this is the first study to pre-grow the bacteria with PM prior to inoculation. All previous studies have either pre-exposed the host to PM or inoculated PM and bacteria together(Liu et al., 2019; Shears et al., 2020; Woo et al., 2018; Yadav et al., 2020). These publications hypothesised that increased bacterial colonisation was due to PM binding to the bacteria thereby promoting transmission throughout the respiratory tract, or PM providing metabolites to support bacterial growth or PM-mediated toxicity damaging epithelial integrity (Liu *et al*., 2019; Yadav *et al*., 2020; Shears *et al*., 2020).

In contrast, our data demonstrate a novel explanation of the detrimental effects of air pollution in that BC directly alters bacterial behaviours to increase colonisation. Supporting this conclusion, the concentration of BC used in our studies does not have a visible effect on host tissue and does not promote *S. aureus* growth (Hussey et al., 2017) and murine colonisation is promoted even when *S. aureus* are pre-grown with BC and BC has not been directly administered to the mice. Together these data demonstrate that major effects of particulates on the host are not essential for increased bacterial colonisation. Similar conclusions come from the work with A549 cells since there was no detectable impact on A549 cell viability and yet *S. aureus* pre-grown in BC prior to infection show a significant increase in staphylococcal adhesion and invasion of these cells.

RNAseq analysis confirms that exposure to BC alters a collection of *S. aureus* genetic responses that have multiple deleterious effects on the host’s ability to combat infection and play important roles in colonisation and dissemination (Pivard et al., 2021). BC increases the transcription of genes for cytotoxins that lyse immune cells (*lukGH, hlgBC, α and β-psms*) (Collins et al., 2020; Tromp & van Strijp, Jos A G, 2020), for factors that inhibit complement (*scin, sbi*) (Sultan *et al*., 2018; Pivard *et al*., 2021) and prevent phagocyte recruitment (chemotaxis-inhibiting protein *chp)*, and are important for *S. aureus* survival in human blood and neutrophils (phospholipase C, *plc*) (White et al., 2014). BC also highly induces expression of the Spl protease genes that play a role in mucin degradation and lung adaptation, with the *splA* mutant showing decreased (Paharik et al., 2016).

Interestingly, BC also induces the SOS response regulators (*lexA, recA*) and effectors (*uvrAB, umuC, hnh, yolD, recJ* and *nrdIFG)*. The SOS response is important for the induced expression of genes important for survival and colonisation of the host including DNA repair, virulence, and immune evasion (Podlesek & Žgur Bertok, 2020). Typically, the SOS response is induced by RecA sensing impairment of bacterial growth and intracellular DNA damage and then initiating the self-cleavage of the LexA repressor protein (Podlesek & Žgur Bertok, 2020), but effect on growth does not seem to be the trigger here because BC does not inhibit *S. aureus* growth and there is no evidence of DNA damage and the transcriptional data do not show other stress responses being activated.

On the contrary, BC represses several genes that are typically induced in response to different stresses, including those involved in oxidative stress (*katA, trxA*), osmotic stress (glycine betaine synthesis *betAB*, proline/betaine transporter *proP*), sulphur metabolism (*cysM, proP, ssuB*) and nitrosative stress (*hmp, ldh*) (Fuchs et al., 2018; Nagarajan & Elasri, 2007). It must be noted though that the negatively regulated genes do not show such a strong uniform response as the genes induced by BC with there being no significant over-representation of down-regulated genes from any functional group.

The BC induction of the toxin, protease, and immune evasion genes (*lukGH, hlgBC, α and β-psms, hla, scin, sbi, chp*) is likely to occur through the activity of the Agr and Sae two-component regulators (Cheung et al., 2011; Geiger et al., 2008) that typically control the expression of these genes. The *S. aureus* Agr quorum sensing system is important for the switch from a colonising state to a more aggressive invasive state through induced expression of toxins and the factors required for dissemination (Jenul & Horswill, 2019). Toxin and immune evasion gene expression is also activated by the Sae regulatory system (Geiger et al., 2008).

Both Agr and Sae have cell membrane located sensors the activity of which can be influenced by a range of different environmental conditions (Geiger et al., 2008; Kavanaugh & Horswill, 2016), although the exact mechanisms involved have not been fully elucidated. It is possible that BC directly interacts with Agr and Sae by either altering environmental signals such as the Agr quorum sensing signal concentrations or activating the membrane-bound sensors to induce gene expression.

The role of Agr and Sae in BC induction of the toxin and immune evasion genes was confirmed by transcriptional analysis of *sae* and *agr* mutants that showed that both Sae and Agr are associated with BC induction of gene expression. The pattern of response differs between the tested genes with either the Agr or Sae regulator or both being required for BC mediated gene regulation. Importantly, BC appears to induce only parts of the Agr and Sae regulons. For example, the expression of adhesin genes that would typically be repressed by Agr (e.g. *spa*) (Cheung et al., 2011) and or induced by Sae (e.g. *fnbA, fnbB, emp*) (Mainiero *et al*., 2010) were not altered in the RNAseq or the qRT-PCR analysis. Therefore, the data suggest that exposure of *S. aureus* to BC prior to or during colonisation of the nares would induce a previously unrecognised regulatory response that increases invasive disease, and which is distinct from previously described patterns of induction of the Agr and Sae regulons.

BC induction of cytotoxins contrasts with the gene regulatory effects observed with other pollutants, e.g. cigarette smoke extract (CSE). As with BC, CSE increases *S. aureus* epithelial cell adhesion and invasion (Kulkarni et al., 2012; Lacoma et al., 2019; McEachern et al., 2015) but in contrast to BC, CSE represses Agr resulting in increased adhesins (Kulkarni et al., 2012) and repressed cytotoxins expression (Lacoma et al., 2019).

Furthermore, our data suggest that BC induces a novel mechanism for increased invasion of epithelial cells. Typically, *S. aureus* invasion of epithelial cells involves Sae-dependent mechanisms involving the fibronectin binding proteins, lipases and toxin induced changes in the cytoskeleton (Josse et al., 2017). The only surface proteins showing induced expression in response to BC in the RNAseq data that are not induced by Agr or Sae are the EfeM/EfeO family lipoprotein (*fepA*) and a Map domain protein both of which have no known role in *S. aureus* invasion. Therefore, the novel mechanism for *S. aureus* invasion induced by BC requires further investigation.

In conclusion, we have provided substantial evidence supporting the novel contention that a single air pollutant, at concentrations non-harmful to bacteria or the host, can specifically alter bacterial behaviour would be expected to have adverse health outcomes. This concept has significant implications for mitigating air pollution toxicity and subsequent adverse health effects, because the currently held hypotheses are restricted to the belief that the toxicity of particle pollutants causes adverse effects by damaging the host directly and that control of pollutant levels need only be limited to concentrations below those that are toxic to humans. This study shows that adverse effects can occur at apparently non-toxic concentrations of pollutant that can alter bacterial behaviour to potentiate infectious disease.

## Experimental procedures

### Bacterial strains and growth conditions

The methicillin resistant *S. aureus* (MRSA) USA300 LAC was used in this study (Kennedy et al., 2008). Transduction with Phage 11 was used to move the *bursa aurealis agrB::*Tn (strain ΦE95) transposon insertion mutation from the Nebraska Transposon Mutant Library (Bae et al., 2008; Fey et al., 2013) and the *saeS*::*Tn917* from strain Newman *sae::Tn917* (Goerke et al., 2001) into USA300 LAC. Mutant strains were confirmed by PCR using gene specific primers (Table S1). Unless otherwise stated, bacteria were grown in Tryptic Soya Broth (TSB; Beckton Dickinson) statically at 37°C in 5% v/v CO_2_.

### Black carbon

Black carbon (BC) (Sigma-Aldrich product number 699632) was dispersed in sterile dH_2_O at 2-10 mg ml^-1^. The particle size of the powder was <500 nm and it contained <500 ppm trace metals. BC is an ideal model particulate for this study as BC does not affect bacterial growth (Hussey, *et al*. 2017), unlike purified synthetic nanoparticles such as Carbon Black which are generally toxic to bacteria (Al-Jumaili et al., 2017).

### Murine colonisation model

Experiments were carried out in accordance with the UK Home Office Project Licence P7B01C07A. Female 8-week-old outbred CD1 mice from Charles River, UK were used. Animals were allowed to acclimatise for 1 week prior to the experiments. Animals were housed in groups of 5, maintained on a 12hr dark/light cycle and allowed unrestricted access to food and water. Prior to use, bacteria were grown in TSB in the presence and absence of 100 μg ml^-1^ BC, to mid exponential phase, and stored in aliquots at -80°C. For use, frozen aliquots were thawed, washed and resuspended in PBS. Mice were intranasally infected with 15 μl containing 1×10^7^ CFU *S. aureus* USA300, or 1×10^7^ CFU LAC mixed with 105 μg of BC as previously described (Hussey et al., 2017). After infection, the mice showed no signs of disease over the following 7 days. At days 1 and 7 post-infection the numbers of bacteria in the nasopharynx, lungs and blood of preselected animals were assessed in homogenised tissue by serial dilution and plating (Hussey et al., 2017). Significance was determined using a Kruskal-Wallis Test with Dunn’s multiple comparison.

### RNA extraction

*S. aureus* strains were grown to late-exponential phase in TSB, with and without 100 μg ml^-1^ BC. To preserve RNA integrity, cultures were treated with RNAprotect (Qiagen) and cells were pelleted and stored at -80°C as per the manufacturer’s instructions. Bacteria were lysed in 200 μl Tris EDTA (TE) buffer containing 100 μg ml^-1^ lysostaphin and 50 μg ml^-1^ proteinase K final concentration. 600 μl of Trizol reagent was added and cells were then mechanically disrupted using a MP Biomedicals FastPrep Instrument and Lysing Matrix B tubes (MP Biomedicals). Black carbon particles were removed by centrifugation at 12, 000 xg and RNA extracted from the bacteria in the supernatant using a Direct-zol RNA Miniprep Plus kit (Zymogen) following the manufacturer’s instructions. Samples were further treated with TURBO DNA-free (Ambion) to ensure complete removal of DNA, which was confirmed via qPCR. RNA concentrations were determined using a Nanodrop spectrophotometer.

### RNAseq

RNA quality and integrity were assessed using a 2100 Bioanalyser and RNA 6000 Nano chip (Agilent), to ensure a minimum RNA integrity value of 8 (Table S4). Samples were depleted for ribosomal RNA and libraries were prepared using ScriptSeq RNA Library Preparation before paired-end sequencing on an Illumina NextSeq550.

RNAseq data quality was assessed using FastQC (v. 0.11.5). Trimmomatic (v. 0.36) was used to remove adaptor sequences, and the read correction tool SOAPec (v. 2.01) was used to identify and repair errors in the read data. The reads were mapped to the *S. aureus* USA300 FRP3757 genome (accession no. CP000255) using HISAT2 (v 2.1.0) and the transcriptome was assembled using STRINGTIE (v. 1.3.3b). The R package DESeq2 was used to test differential gene expression between the samples, and gene expression is expressed as the Log2 Fold Change (L2FC) in expression relative to growth without BC. The screening threshold for the results was set at >1 or <-1 L2FC using an adjusted p-Value (pADJ) of 0.001.

To determine whether any functional groups of genes were significantly up-regulated or down-regulated in response to BC, GO (gene ontology) Enrichment Analysis was carried out and the Fisher’s exact test with a p value ≤ 0.05 was used to test the enrichment in each category (Ashburner et al., 2000; Gene, 2021; Mi et al., 2019). Additional gene function data, including TIGRFAM functional groups, were extracted for each locus from the AureoWiki database, which provides a pan-genome approach to functional annotation of genes (Fuchs et al., 2018).

### Quantitative reverse transcriptase PCR (qRT-PCR)

Total RNA converted into cDNA using Superscript IV VILO Master Mix reverse transcriptase (Invitrogen), and 0.5 ng of cDNA was used for each qPCR reaction. qRT-PCR was done using SYBR Green Master mix (Applied Biosystems) in a 7300 Fast System (Applied Biosystems) following the manufacturer’s instructions. Relative gene expression for each of the sample genes (for primer details see Table S1) was normalized to the expression of the endogenous control genes *gyrB* and 16S rRNA and expressed relative to the LAC wildtype strain cultured without BC, using the ΔΔCt method to calculate RQ (2^-ΔΔCt^) (Livak & Schmittgen, 2001). Significance was determined by a Kruskal Wallis Test with Dunn’s multiple comparison test (* = p<0.05, **= p<0.01).

### Epithelial cell adhesion, invasion and persistence

For bacterial adhesion and invasion studies, the human Type II-like bronchial epithelial cell line A549 was used. 24-well tissue culture plates were seeded with 1×10^5^ cells in RPMI with 1% v/v foetal bovine calf serum (FBS) and grown to 70-100% confluency prior to inoculation with *S. aureus*. A549 cells were inoculated with 1× 10^7^ CFU of *S. aureus* under the following conditions: (i) *S. aureus* LAC alone, (ii) *S. aureus* LAC plus 100 μg ml^-1^ of BC or (iii) *S. aureus* LAC grown in the presence of 100 μg ml^-1^ BC with particles diluted from inoculum All doses were confirmed by serial dilution and plating. Infected cells were incubated at 37°C in 5% v/v CO_2_ for 2 hours. For adhesion assays, cells were washed in PBS and lysed in 1% v/v Triton-X-100 for 10 minutes. Bacterial CFU was determined by serial dilution and plating. For invasion and persistence assays, 2 hours post-inoculation A549 cells were washed and resuspended in RPMI containing 300 μg ml^-1^ gentamicin for 2 hours (invasion) before washing and lysing the cells as described above, followed by serial dilution and plating to determine CFU (Richards et al., 2015). Data are presented as the mean of at least 3 independent biological replicates (+/- SEM) and significance was determined by One-way ANOVA with Tukey’s multiple comparison test.

### Cytotoxicity

A549 cytotoxicity was measured by the presence of lactate dehydrogenase (LDH) in the culture medium. LDH release was measured using CyQuant LDH Cytotoxicity Assay Kit (Invitrogen) as per the manufacturer’s instructions. LDH activity was assayed in supernatant from uninfected cells (spontaneous damage), cells infected with bacteria alone and with 100 μg/ml BC and cells exposed to 100 μg/ml BC alone (to determine cell damage from BC specifically). Absorbance was measured at 490 nm and 680 nm (background) and the background value was subtracted from the 490 nm reading to give LDH activity. % cytotoxicity was calculated as follows: ((Supernatant LDH activity – Spontaneous LDH activity) / (MAX LDH activity – Spontaneous LDH activity)) x 100

Data are presented as the mean of 3 independent biological replicates (+/- SEM) and significance was determined by one-way ANOVA with Tukey’s multiple comparison test.

## Acknowledgements

USA300 JE2 and JE2 *agr*::Tn strains were obtained through the Network on Antimicrobial Resistance in *Staphylococcus aureus* (NARSA) Program: USA300 supported under NIAID/ NIH Contract No. HHSN272200700055C. We would like to thank Andrew Briscoe at the Core Research Laboratories, Natural History Museum for the RNAseq library preparation and sequencing. JP and SJKH were supported by a Leverhulme Trust grant (RPG-2015-183) awarded to JAM, PWA, JMK, PSM; LC was supported by a National Centre for Atmospheric Science Air Pollution Science Training Studentship Programme. LP was supported by MRC DTP IMPACT studentship.

