## Supplemental Tables for "Air pollution induces *Staphylococcus aureus* USA300 respiratory tract colonisation mediated by specific bacterial genetic responses dependent on the global virulence gene regulators Agr and Sae"

Supplementary Tables:

**Table S1. Primers used in this study**.

| **Name** | **Gene** | **Nucleotide sequence (5’ to 3’)^a^** | **Tm** | **PCR efficiency** |
| --- | --- | --- | --- | --- |
| gyrB-F  gyrB-R | *gyrB* | GACTGATGCCGATGTGGA  AACGGTGGCTGTGCAATA | 60 | 95.8% |
| splF_qRT_F  splF_qRT_R | *splF* | CCTGGTAGCTCTGGTTCACC  CCTGATGGCTTATTACCGGC | 60 | 93.9% |
| CHIPS_qRT_F CHIPS_qRT_R | *chp* | GCTTTTACTTTTGAACCGTTTCC  CGTTCATCTAATTTTCCTAGCGT | 60 | 88.9% |
| betB_qRT_F  betB_qRT_R | *betB* | GCAGATAAAGACGGTGGCGA  GTGAACAACCCGTAGCAAGC | 60 | 96.3% |
| ecb_qRT_F  ebc_qRT_R | *ecb* | TTGCCGGTGAATCTCATGCA  GTGCTTTACGGTGTGTTGCA | 60 | 94.7% |
| epiA_qRT_F  epiA_qRT_R | *epiA* | AACTCAAATGATTCAGCAGGTGA  AAACTACCAGTCTTAGCACAACC | 60 | 97.9% |
| hla_qRT_F  hla_qRT_R | *hla* | CTTGGAACCCGGTATATGGCA  GCGAAGTCTGGTGAAAACCC | 60 | 89.9% |
| clfA_qRT_F  clfA_qRT_R | *clfA* | AGTGCGCCTAGAATGAGAGC  GCGGATACACAGTCGTACCA | 60 | 98.8% |
| fnbA_qRT_F  fnbA_qRT_R | *fnbA* | GCGAAAGTGGAAACGGGTAC  TAGTACCGCTCGTTGTCCTG | 60 | 92.2% |
| fnbB_qRT_F  fnbB_qRT_R | *fnbB* | CACAATCATACAGCGCGACA  TGGTGCTTGCACAGTTTTCG | 60 | 98.4% |
| emp_qRT_F  emp_qRT_R | *emp* | CAGAATCGCCTAGATATACACATCCA  GCATGCCCTGGTGTAACAAAATT | 60 | 87% |
| katA_qRT_F  katA_qRT_R | *katA* | GGAGCGTGACATTCGAGGAT  AGGATCTCGTTTCACCGCAC | 60 | 90.3% |
| sodA_qRT_F  sodA_qRT_R | *soda* | CCAATGTAGTCAGGGCGTTTG  AGTCGTAAACAATGGCCAGT | 60 | 90.5% |
| PSMB_qRT_F  PSMB_qRT_R | *PSMB* | CAAGCTGCACAACAACATGA  ACCTAGTAAACCCACACCGT | 60 | 101.8% |
| lukS_qRT_F  lukS_qRT_R | *lukS* | GTCTCAAAACAAATGACCCCAA  ACCTCCTGTTGATGGACCAC | 60 | 89.6% |
| kdpD_qRT_F  kdpD_qRT_R | *kdpD* | TGATTCGGACTTTAAGCGGACA  CCTCCACCATTCGTACCAAGA | 60 | 90.2% |

Table S2. **Genes significantly upregulated at a L2FC >1in response to BC, grouped into TIGRFAM functional groups (Main). * entries do not have an official TIGRFAM entry and have been annotated based on literature review of their functions**

| Locus Tag | Gene name | Product Description | L2FC | Fold Change | pAdj | TIGRFAM No. | Main Role | Sub Role |
| --- | --- | --- | --- | --- | --- | --- | --- | --- |
| SAUSA300_RS02340 | *metQ2* | dipeptide ABC transporter glycylmethionine-binding lipoprotein | 1.40 | 2.63 | 2.67E-24 | TIGR00363 | Cell envelope | - |
| SAUSA300_RS10495 |  | MAP domain-containing protein | 1.20 | 2.30 | 2.10E-12 | - | *Cellular processes | *Adhesion to cells |
| SAUSA300_RS14555 | *-* | S-adenosyl-l-methionine hydroxide adenosyltransferase family protein | 1.09 | 2.13 | 1.31E-04 | TIGR04507 | Cellular processes | Biosynthesis of natural products |
| SAUSA300_RS13080 | *hlgB* | bi-component gamma-hemolysin HlgAB/HlgCB subunit B | 1.52 | 2.87 | 3.28E-18 | TIGR01002 | Cellular processes | Toxin production and resistance |
| SAUSA300_RS00515 | *plc* | phosphatidylinositol-specific phospholipase C | 1.42 | 2.68 | 2.02E-27 | - | *Cellular processes | *Pathogenesis |
| SAUSA300_RS13075 | *hlgC* | bi-component gamma-hemolysin HlgCB subunit C | 1.24 | 2.36 | 1.83E-07 | TIGR01002 | Cellular processes | Toxin production and resistance |
| SAUSA300_RS10845 | *lukG* | bi-component leukocidin LukGH subunit G | 1.19 | 2.29 | 2.65E-15 | TIGR01002 | Cellular processes | Toxin production and resistance |
| SAUSA300_RS10505 | *hlb-1* | sphingomyelin phosphodiesterase | 1.10 | 2.14 | 7.39E-11 | TIGR03395 | Cellular processes | Pathogenesis |
| SAUSA300_RS07540 |  | Panton-Valentine bi-component leukocidin subunit F | 1.06 | 2.09 | 8.48E-09 | TIGR01002 | Cellular processes | Toxin production and resistance |
| SAUSA300_RS10850 | *lukH* | bi-component leukocidin LukGH subunit H | 1.02 | 2.03 | 8.54E-08 | TIGR01002 | Cellular processes | Toxin production and resistance |
| SAUSA300_RS05720 | *hla* | alpha-hemolysin | 1.93 | 3.81 | 1.42E-44 | TIGR01002 | Cellular processes | Toxin production and resistance |
| SAUSA300_RS01705 | *gehB* | YSIRK domain-containing triacylglycerol lipase Lip2/Geh | 1.54 | 2.90 | 1.22E-24 | TIGR01168 | Cellular processes | - |
| SAUSA300_RS06840 | *mucB* | DNA repair protein MucB | 1.48 | 2.78 | 1.11E-18 | - | *DNA metabolism | *DNA replication, recombination, and repair |
| SAUSA300_RS06370 | *recA* | DNA recombination/repair protein RecA | 1.38 | 2.61 | 4.59E-24 | TIGR02012 | DNA metabolism | DNA replication, recombination, and repair |
| SAUSA300_RS04005 | *uvrA* | excinuclease ABC subunit UvrA | 1.38 | 2.61 | 6.91E-25 | TIGR00630 | DNA metabolism | DNA replication, recombination, and repair |
| SAUSA300_RS04985 | *comK1* | competence protein ComK | 1.28 | 2.42 | 1.24E-04 | - | *DNA metabolism | *DNA interactions |
| SAUSA300_RS04000 | *uvrB* | excinuclease ABC subunit UvrB | 1.09 | 2.13 | 8.22E-19 | TIGR00631 | DNA metabolism | DNA replication, recombination, and repair |
| SAUSA300_RS10420 | *-* | YolD-like family protein | 1.07 | 2.10 | 3.75E-06 | - | *DNA metabolism | - |
| SAUSA300_RS08675 | *recJ* | single-stranded-DNA-specific exonuclease RecJ | 1.03 | 2.04 | 1.71E-13 | TIGR00644 | DNA metabolism | DNA replication, recombination, and repair |
| SAUSA300_RS06750 | *sbcC* | SMC family ATPase | 1.03 | 2.04 | 1.86E-08 | TIGR00618 | DNA metabolism | DNA replication, recombination, and repair |
| SAUSA300_RS06710 | *lexA* | transcriptional repressor LexA | 1.08 | 2.11 | 1.51E-17 | TIGR00498 | DNA metabolism; Regulatory functions | DNA interactions; DNA replication, recombination, and repair |
| SAUSA300_RS06445 | *glpD* | glycerol-3-phosphate dehydrogenase/oxidase | 1.57 | 2.96 | 4.52E-25 | TIGR03377 | Energy metabolism | Anaerobic |
| SAUSA300_RS10530 | *chp* | chemotaxis-inhibiting protein CHIPS | 2.25 | 4.75 | 1.31E-34 | - | *Immune evasion | *Pathogenesis |
| SAUSA300_RS10525 | *scn* | complement inhibitor SCIN-A | 1.85 | 3.60 | 1.11E-56 | TIGR04561 | Immune evasion | Pathogenesis |
| SAUSA300_RS15090 | *-* | phenol-soluble modulin PSM-alpha-3 | 1.82 | 3.53 | 1.59E-38 | - | *Immune evasion | *Toxin production and resistance |
| SAUSA300_RS15730 | *-* | phenol-soluble modulin PSM-alpha-4 | 1.81 | 3.50 | 1.25E-24 | - | *Immune evasion | *Toxin production and resistance |
| SAUSA300_RS05795 | *psm_2* | beta-class phenol-soluble modulin | 1.78 | 3.45 | 5.79E-18 | - | *Immune evasion | *Toxin production and resistance |
| SAUSA300_RS15735 | *-* | phenol-soluble modulin PSM-alpha-2 | 1.78 | 3.44 | 6.10E-30 | - | *Immune evasion | *Toxin production and resistance |
| SAUSA300_RS15740 | *-* | phenol-soluble modulin PSM-alpha-1 | 1.62 | 3.07 | 1.69E-15 | - | *Immune evasion | *Toxin production and resistance |
| SAUSA300_RS05790 | *psm_1* | beta-class phenol-soluble modulin | 1.42 | 2.68 | 3.86E-16 | - | *Immune evasion | *Toxin production and resistance |
| SAUSA300_RS13060 | *sbi* | immunoglobulin-binding protein Sbi | 1.08 | 2.11 | 1.26E-17 | - | *Immune evasion | *Toxin production and resistance |
| SAUSA300_RS04395 | *ear* | DUF4888 domain-containing protein | 1.36 | 2.57 | 4.32E-18 | TIGR01558 | Mobile and extrachromosomal element functions | Prophage functions |
| SAUSA300_RS10660 | *-* | HNH endonuclease | 1.22 | 2.34 | 1.25E-04 | TIGR03508 | Mobile and extrachromosomal element functions | Prophage functions |
| SAUSA300_RS09595 | *splF* | serine protease SplF | 1.99 | 3.99 | 5.13E-16 | TIGR02037 | Protein fate | Degradation of proteins, peptides, and glycopeptides; Protein folding and stabilization |
| SAUSA300_RS09605 | *splD* | serine protease SplD | 1.91 | 3.75 | 1.81E-13 | TIGR02037 | Protein fate | Degradation of proteins, peptides, and glycopeptides; Protein folding and stabilization |
| SAUSA300_RS09600 | *splE* | serine protease SplE | 1.50 | 2.83 | 1.37E-06 | TIGR02037 | Protein fate | Degradation of proteins, peptides, and glycopeptides; Protein folding and stabilization |
| SAUSA300_RS14170 | *nrdG* | anaerobic ribonucleoside-triphosphate reductase activating protein | 1.30 | 2.46 | 5.77E-27 | TIGR02491 | Protein fate; Purines, pyrimidines, nucleosides, and nucleotides | 2'-Deoxyribonucleotide metabolism; Protein modification and repair |
| SAUSA300_RS09620 | *splA* | serine protease SplA | 2.29 | 4.88 | 3.02E-25 | TIGR02038 | Protein fate; Regulatory functions | Degradation of proteins, peptides, and glycopeptides; Protein interactions |
| SAUSA300_RS09610 | *splC* | serine protease SplC | 2.14 | 4.40 | 1.06E-17 | TIGR02038 | Protein fate; Regulatory functions | Degradation of proteins, peptides, and glycopeptides; Protein interactions |
| SAUSA300_RS09615 | *splB* | serine protease SplB | 2.02 | 4.04 | 1.32E-14 | TIGR02038 | Protein fate; Regulatory functions | Degradation of proteins, peptides, and glycopeptides; Protein interactions |
| SAUSA300_RS08875 | *infC* | translation initiation factor IF-3 | 1.06 | 2.09 | 2.19E-18 | TIGR00168 | Protein synthesis | Translation factors |
| SAUSA300_RS03850 | *nrdF* | class 1b ribonucleoside-diphosphate reductase subunit beta | 1.04 | 2.05 | 5.26E-12 | TIGR04171 | Purines, pyrimidines, nucleosides, and nucleotides | 2'-Deoxyribonucleotide metabolism |
| SAUSA300_RS03840 | *nrdI* | class Ib ribonucleoside-diphosphate reductase assembly flavoprotein NrdI | 1.04 | 2.05 | 7.34E-13 | TIGR00333 | Purines, pyrimidines, nucleosides, and nucleotides | 2'-Deoxyribonucleotide metabolism |
| SAUSA300_RS11200 | *kdpD* | sensor histidine kinase KdpD | 1.03 | 2.04 | 3.99E-11 | TIGR02966 | Signal transduction | Two-component systems |
| SAUSA300_RS11190 | *kdpA* | potassium-transporting ATPase subunit A | 2.65 | 6.30 | 1.28E-04 | TIGR00680 | Transport and binding proteins | Cations and iron carrying compounds |
| SAUSA300_RS02335 | *metP2* | ABC transporter permease | 1.34 | 2.53 | 7.92E-17 | TIGR01097 | Transport and binding proteins | Anions |
| SAUSA300_RS01825 | *fepA* | EfeM/EfeO family lipoprotein | 1.18 | 2.27 | 7.97E-15 | TIGR04358 | Transport and binding proteins | Cations and iron carrying compounds |
| SAUSA300_RS02330 | *metN2* | methionine ABC transporter ATP-binding protein | 1.10 | 2.15 | 2.63E-16 | TIGR02314 | Transport and binding proteins | Amino acids and amines |
| SAUSA300_RS00185 | *-* | DUF1643 domain-containing protein | 1.72 | 3.29 | 8.60E-04 | - | - | - |
| SAUSA300_RS06715 |  | hypothetical protein | 1.52 | 2.86 | 1.32E-11 | - | - | - |
| SAUSA300_RS01835 | *fepC* | iron permease FTR1 family protein | 1.39 | 2.61 | 2.22E-16 | TIGR00145 | - | General |
| SAUSA300_RS04990 | *-* | IDEAL domain-containing protein | 1.05 | 2.07 | 2.75E-05 | - | - | - |

Table S3. **Genes significantly downregulted at a L2FC <-1 in response to BC, grouped into TIGRFAM functional groups (Main) * entries do not have an official TIGRFAM entry and have been annotated based on literature review of their functions**

| Locus Tag | Gene Name | Product Description | L2FC | Fold Change | pAdj | TIGRFAM number | Main Role | Sub Role |
| --- | --- | --- | --- | --- | --- | --- | --- | --- |
| SAUSA300_RS05755 | *argF* | ornithine carbamoyltransferase | -1.82 | -3.53 | 2.65E-10 | TIGR00658 | Amino acid biosynthesis | Glutamate family |
| SAUSA300_RS02320 | *mccA* | cysteine synthase family protein | -1.65 | -3.14 | 8.64E-31 | TIGR01136 | Amino acid biosynthesis | Serine family |
| SAUSA300_RS13660 | *ddh* | D-lactate dehydrogenase | -1.66 | -3.15 | 1.39E-51 | TIGR01327 | Amino acid biosynthesis | Serine family |
| SAUSA300_RS02030 | *nfrA* | NADPH-dependent oxidoreductase | -1.14 | -2.21 | 6.76E-11 | TIGR02476 | Biosynthesis of cofactors, prosthetic groups, and carriers | Heme, porphyrin, and cobalamin |
| SAUSA300_RS13655 | *frp* | NAD(P)H-dependent oxidoreductase | -1.26 | -2.40 | 4.71E-20 | TIGR02476 | Biosynthesis of cofactors, prosthetic groups, and carriers | Heme, porphyrin, and cobalamin |
| SAUSA300_RS10360 | *nadE* | ammonia-dependent NAD(+) synthetase | -1.09 | -2.13 | 2.54E-17 | TIGR00552 | Biosynthesis of cofactors, prosthetic groups, and carriers | Pyridine nucleotides |
| SAUSA300_RS02700 | *pdxS* | pyridoxal 5'-phosphate synthase lyase subunit PdxS | -1.74 | -3.35 | 2.91E-22 | TIGR00343 | Biosynthesis of cofactors, prosthetic groups, and carriers | Pyridoxine |
| SAUSA300_RS02705 | *pdxT* | pyridoxal 5'-phosphate synthase glutaminase subunit PdxT | -1.87 | -3.66 | 6.23E-46 | TIGR03800 | Biosynthesis of cofactors, prosthetic groups, and carriers | Pyridoxine |
| SAUSA300_RS14130 | *betA* | oxygen-dependent choline dehydrogenase | -1.74 | -3.34 | 2.34E-10 | TIGR01810 | Cellular processes | Adaptations to atypical conditions |
| SAUSA300_RS14135 | *betB* | betaine-aldehyde dehydrogenase | -2.80 | -6.96 | 2.35E-16 | TIGR01804 | Cellular processes | Adaptations to atypical conditions |
| SAUSA300_RS11395 | *ptpB* | low molecular weight protein arginine phosphatase | -1.81 | -3.51 | 4.04E-29 | TIGR02691 | Cellular processes | Detoxification |
| SAUSA300_RS01460 | *-* | ABC transporter permease | -1.41 | -2.66 | 1.49E-14 | TIGR03061 | Cellular processes | Enzymes of unknown specificity |
| SAUSA300_RS07865 | *-* | DUF1672 domain-containing protein | -1.26 | -2.40 | 1.47E-09 | TIGR02898 | Cellular processes | Sporulation and germination |
| SAUSA300_RS09670 | *epiA* | gallidermin/nisin family lantibiotic | -1.92 | -3.78 | 1.88E-05 | TIGR03731 | Cellular processes | Toxin production and resistance |
| SAUSA300_RS14615 | *-* | arylamine N-acetyltransferase | -1.22 | -2.34 | 5.73E-10 | - | *Cellular processes; Central intermediary metabolism | - |
| SAUSA300_RS02920 | *sdrD* | MSCRAMM family adhesin SdrD | -1.41 | -2.66 | 2.82E-20 | TIGR01168 | Cellular processes; Transport and binding proteins | Pathogenesis |
| SAUSA300_RS00995 | *-* | alpha-keto acid decarboxylase family protein | -1.85 | -3.59 | 1.14E-27 | TIGR03393 | Central intermediary metabolism | Other |
| SAUSA300_RS09050 | *-* | class I SAM-dependent methyltransferase | -1.03 | -2.04 | 9.50E-07 | TIGR00497 | DNA metabolism | Restriction/modification |
| SAUSA300_RS05760 | *arcC1* | carbamate kinase | -1.48 | -2.78 | 3.14E-05 | TIGR00746 | Energy metabolism | Amino acids and amines |
| SAUSA300_RS00905 | *-* | hypothetical protein | -1.63 | -3.09 | 6.92E-05 | TIGR03908 | Energy metabolism | Amino acids and amines |
| SAUSA300_RS12320 | *fdhD* | formate dehydrogenase accessory sulfurtransferase FdhD | -1.03 | -2.04 | 1.46E-05 | TIGR00129 | Energy metabolism | Anaerobic; Electron transport |
| SAUSA300_RS00990 | *-* | isochorismatase family protein | -1.02 | -2.02 | 3.79E-04 | TIGR03614 | Energy metabolism | Carbohydrates, organic alcohols, and acids |
| SAUSA300_RS13045 | *gpmA* | phosphoglycerate mutase | -1.16 | -2.24 | 1.55E-16 | TIGR01258 | Energy metabolism | Carbohydrates, organic alcohols, and acids |
| SAUSA300_RS05620 | *trxA* | thioredoxin | -1.05 | -2.07 | 3.18E-04 | TIGR01068 | Energy metabolism | Electron transport |
| SAUSA300_RS01240 | *hmp* | nitric oxide dioxygenase | -1.00 | -2.00 | 1.96E-13 | TIGR02160 | Energy metabolism | Other |
| SAUSA300_RS02970 | *-* | HAD family hydrolase | -1.06 | -2.08 | 7.48E-04 | TIGR01449 | Energy metabolism | Sugars |
| SAUSA300_RS06680 | *katA* | catalase | -1.49 | -2.80 | 7.35E-22 | - | *Immune evasion | *Adaptations to atypical conditions |
| SAUSA300_RS14005 | *-* | TIGR04197 family type VII secretion effector | -1.94 | -3.83 | 4.35E-04 | TIGR04197 | Immune evasion | - |
| SAUSA300_RS01490 | *esxA* | WXG100 family type VII secretion effector EsxA | -1.31 | -2.47 | 9.84E-29 | TIGR03930 | Immune evasion | Pathogenesis |
| SAUSA300_RS00560 | *-* | oleate hydratase | -1.33 | -2.51 | 1.18E-30 | TIGR02732 | Immune evasion | Pathogenesis |
| SAUSA300_RS06940 | *-* | oligoendopeptidase F | -1.10 | -2.15 | 7.22E-16 | TIGR00181 | Protein fate | Degradation of proteins, peptides, and glycopeptides |
| SAUSA300_RS04855 | *pepF* | oligoendopeptidase F | -1.14 | -2.21 | 4.46E-24 | TIGR00181 | Protein fate | Degradation of proteins, peptides, and glycopeptides |
| SAUSA300_RS05990 | *-* | hypothetical protein | -1.74 | -3.34 | 7.62E-17 | TIGR03302 | Protein fate | Protein and peptide secretion and trafficking |
| SAUSA300_RS04995 | *lplA1* | lipoate--protein ligase | -1.04 | -2.05 | 5.43E-11 | TIGR00545 | Protein fate | Protein modification and repair |
| SAUSA300_RS12640 | *-* | sodium ABC transporter permease | -1.38 | -2.61 | 6.74E-26 | TIGR01302 | Purines, pyrimidines, nucleosides, and nucleotides | Purine ribonucleotide biosynthesis |
| SAUSA300_RS11905 | *-* | MerR family transcriptional regulator | -1.48 | -2.78 | 6.78E-09 | TIGR02044 | Regulatory functions | DNA interactions |
| SAUSA300_RS05670 | *ecb* | complement convertase inhibitor Ecb | -1.91 | -3.75 | 9.19E-13 | - | *Transport and binding proteins | *Adhesion to cells |
| SAUSA300_RS05690 | *efb* | fibrinogen-binding protein | -1.46 | -2.76 | 1.40E-18 | - | *Transport and binding proteins | *Adhesion to cells |
| SAUSA300_RS00915 | *ssuB* | ABC transporter ATP-binding protein | -1.10 | -2.15 | 5.54E-18 | TIGR01184 | Transport and binding proteins | Anions; Other |
| SAUSA300_RS01455 | *-* | ABC transporter ATP-binding protein | -1.15 | -2.22 | 2.18E-05 | TIGR03864 | Transport and binding proteins | Carbohydrates, organic alcohols, and acids |
| SAUSA300_RS00665 | *-* | MFS transporter | -1.49 | -2.81 | 2.97E-07 | TIGR03025 | Transport and binding proteins | Enzymes of unknown specificity |
| SAUSA300_RS03315 | *mntC* | metal ABC transporter substrate-binding protein | -1.10 | -2.14 | 5.06E-24 | TIGR03772 | Transport and binding proteins | Unknown substrate |
| SAUSA300_RS02975 | *proP* | proline/betaine transporter | -1.21 | -2.31 | 1.01E-17 | TIGR00883 | Transport and binding proteins | Unknown substrate |
| SAUSA300_RS11450 | *-* | DUF2529 domain-containing protein | -1.10 | -2.14 | 2.24E-04 | - | - | - |
| SAUSA300_RS02100 | *-* | DUF1304 domain-containing protein | -1.01 | -2.01 | 7.88E-05 | - | - | - |
| SAUSA300_RS13480 |  | hypothetical protein | -1.05 | -2.06 | 1.84E-06 | - | - | - |
| SAUSA300_RS06380 | *-* | hypothetical protein | -1.06 | -2.09 | 2.60E-04 | - | - | - |
| SAUSA300_RS06665 | *-* | hypothetical protein | -1.07 | -2.10 | 9.58E-05 | - | - | - |
| SAUSA300_RS01620 | *-* | DUF4064 domain-containing protein | -1.10 | -2.15 | 4.36E-14 | - | - | - |
| SAUSA300_RS00910 | *-* | DUF4242 domain-containing protein | -1.11 | -2.16 | 9.03E-22 | - | - | - |
| SAUSA300_RS01230 | *-* | DUF488 domain-containing protein | -1.17 | -2.25 | 2.27E-13 | - | - | - |
| SAUSA300_RS01485 | *-* | CHAP domain-containing protein | -1.21 | -2.31 | 9.33E-14 | - | - | - |
| SAUSA300_RS14315 | *isaB* | immunodominant staphylococcal antigen IsaB | -1.24 | -2.36 | 4.53E-14 | - | - | - |
| SAUSA300_RS04765 | *-* | DUF2929 domain-containing protein | -1.32 | -2.50 | 4.53E-05 | - | - | - |
| SAUSA300_RS11230 | *-* | hypothetical protein | -1.38 | -2.60 | 9.03E-09 | - | - | - |
| SAUSA300_RS13140 |  | type I toxin-antitoxin system Fst family toxin | -1.39 | -2.62 | 8.03E-05 | - | - | - |
| SAUSA300_RS14030 | *-* | glyoxalase/bleomycin resistance/extradiol dioxygenase family protein | -1.45 | -2.73 | 1.12E-06 | - | - | - |
| SAUSA300_RS05495 | *-* | YlbG family protein | -1.52 | -2.87 | 1.25E-04 | - | - | - |
| SAUSA300_RS12650 | *-* | DUF805 domain-containing protein | -1.66 | -3.15 | 2.39E-13 | TIGR02353 | - | - |
| SAUSA300_RS01470 | *-* | hypothetical protein | -2.15 | -4.43 | 4.80E-27 | - | - | - |
| SAUSA300_RS02105 | *-* | SDR family oxidoreductase | -1.10 | -2.14 | 3.16E-10 | TIGR03649 | - | Enzymes of unknown specificity |
| SAUSA300_RS08610 | *-* | LLM class flavin-dependent oxidoreductase | -1.12 | -2.17 | 1.02E-12 | TIGR03558 | - | Enzymes of unknown specificity |
| SAUSA300_RS11900 | *-* | aldo/keto reductase | -1.56 | -2.95 | 3.83E-25 | TIGR01293 | - | Enzymes of unknown specificity |

**Table S4**

RNA integrity and concentrations of samples sent for RNAseq.

| **Bacterial Strain** | **Black Carbon** | **Replicate** | **RIN (RNA integrity number)** | **Concentration (ng/µl)** |
| --- | --- | --- | --- | --- |
| USA300 LAC | Control | 1 | 9.2 | 607 |
| USA300 LAC | 100 µg/ml | 1 | 9.5 | 653 |
| USA300 LAC | Control | 2 | 9.4 | 787 |
| USA300 LAC | 100 µg/ml | 2 | 9.5 | 722 |
